# Exploring the role of Polycomb recruitment in *Xist*-mediated silencing of the X chromosome in ES cells

**DOI:** 10.1101/495739

**Authors:** Aurélie Bousard, Ana Cláudia Raposo, Jan Jakub Żylicz, Christel Picard, Vanessa Borges Pires, Yanyan Qi, Laurène Syx, Howard Y. Chang, Edith Heard, Simão Teixeira da Rocha

## Abstract

*Xist* RNA has been established as the master regulator of X-chromosome inactivation (XCI) in female eutherian mammals but its mechanism of action remains unclear. By creating novel *Xist* mutants at the endogenous locus in mouse embryonic stem (ES) cells, we dissect the role of the conserved A-B-C-F repeats. We find that transcriptional silencing can be largely uncoupled from Polycomb repressive complex 1 and 2 (PRC1/2) recruitment, which requires repeats B and C. *Xist* ΔB+C RNA specifically loses interaction with PCGF3/5 subunits of PRC1, while binding of other *Xist* partners is largely unaffected. However, a slight relaxation of transcriptional silencing in *Xist* ΔB+C indicates a role for PRC1/2 proteins in early stabilization of gene repression. Distinct modules within the *Xist* RNA are therefore involved in the convergence of independent chromatin modification and gene repression pathways. In this context, Polycomb recruitment seems to be of moderate relevance in the initiation of silencing.

## Introduction

Long non-coding RNAs (lncRNAs) are a class of non-protein coding RNAs of > 200 nucleotides, that are frequently capped, spliced and polyadenylated. Some are located in the nucleus and have been implicated in transcriptional regulation and recruitment of chromatin modifiers, using still poorly defined molecular mechanisms [reviewed in (Rutenberg-Schoenberg et al., 2016; Schmitz et al., 2016)]. *Xist* (*X-inactive-specific transcript*) lncRNA represents the most studied paradigm of a nuclear RNA with documented roles in transcription regulation and recruitment of chromatin modifiers in female eutherian mammals [reviewed in (da Rocha and Heard, 2017)]. *Xist* lncRNA is ultimately expressed from only one of the two X chromosomes, “coating” in *cis* its chromosome territory and triggering a cascade of events that result in chromosome-wide gene silencing and formation of facultative heterochromatin [reviewed in (da Rocha and Heard, 2017)]. How *Xist* coordinates these two processes, and their causal relationship, is still unclear. In this context, the Polycomb group (PcG) proteins, that modify chromatin at early stages of XCI are of particular interest [reviewed in (Escamilla-Del-Arenal et al., 2011)].

Recruitment of PcG proteins following *Xist* RNA coating is an early event during XCI. Both PRC2 and PRC1 are recruited to lay down H3K27me3 and H2AK119ub on the future inactive X chromosome (Xi), respectively (de Napoles et al., 2004; Plath et al., 2003; Silva et al., 2003). Both canonical PRC1, with a CBX7 subunit, and non-canonical versions, with RYBP/YAF2 subunits, are known to be recruited to the Xi (Almeida et al., 2017; Leeb and Wutz, 2007; Tavares et al., 2012). Previous studies showed that *Xist* RNA indeed interacts with non-canonical PRC1 components, as well as with the RNA binding protein, hnRNPK (Chu et al., 2015). Recently, it was discovered that the non-canonical PCGF3/5-PRC1, associates with *Xist* RNA via hnRNPK and appears to mediate early H2AK119ub deposition. This may then be required for PRC2 recruitment (Almeida et al., 2017). Consistent with this, our previous work demonstrated that PRC2 recruitment involves its cofactor JARID2 (da Rocha et al., 2014). This in turn may be recruited to the Xi via binding to the PRC1-mark H2AK119ub (Cooper et al., 2016).

The role of PRC2 in XCI was first uncovered through the analysis of a mouse hypomorph knock-out for *Eed*, a PRC2 component, in which loss in maintenance of XCI was seen in the extra-embryonic tissues of female embryos (Kalantry et al., 2006; Wang et al., 2001). In contrast, the role of PRC2 or PRC1 during XCI in embryonic lineages or in differentiating ES cells, remains inconclusive with slightly inconsistent results (Almeida et al., 2017; Kalantry and Magnuson, 2006; Leeb and Wutz, 2007; Silva et al., 2003). These discrepancies arise in part from the models used to address this question. *In vivo* analysis of XCI has been confounded by severe developmental abnormalities and early lethality upon disruption of PRC2 function (Kalantry and Magnuson, 2006; Silva et al., 2003). Similarly, *ex vivo* analyses usually involved *Xist* transgenes on autosomes which may not fully recapitulate the chromatin requirements of the X chromosome during XCI (Almeida et al., 2017; Leeb and Wutz, 2007; Loda et al., 2017). It is in this context that we set out to address PcG function in *Xist*-dependent transcriptional silencing during ES cell differentiation.

*Xist* is an unusually long RNA (15,000-17,000 nt) with low overall sequence conservation, except for a series of unique tandem repeats, named A-to-F (Figure 1A) (Brown, 1991; Nesterova et al., 2001; Yen et al., 2007). The most conserved and best studied is the A repeat, which is essential for *Xist*-mediated gene silencing (Wutz et al., 2002). The A repeat interacts specifically with proteins such as SPEN and RBM15 both believed to be involved in its gene silencing role (Chu et al., 2015; Lu et al., 2016; McHugh et al., 2015; Moindrot et al., 2015; Monfort et al., 2015; Patil et al., 2016). Other *Xist* RNA repeat regions have been implicated in the recruitment of factors involved in *cis*-localization (*e.g.*, recruitment of CIZ1 matrix attachment protein by the E repeat) (Ridings-Figueroa et al., 2017; Sunwoo et al., 2017) or Polycomb chromatin modifications (da Rocha et al., 2014; Pintacuda et al., 2017). We previously showed that a region spanning F, B and C repeats is critical for PRC2 recruitment to the Xi (da Rocha et al., 2014). More recently, using an autosomal *Xist* transgene, it was reported that a 600 bp *Xist* region containing the B repeat was necessary for PRC2 and PRC1 recruitment through direct binding of hnRNPK (Pintacuda et al., 2017).

**Figure 1.**
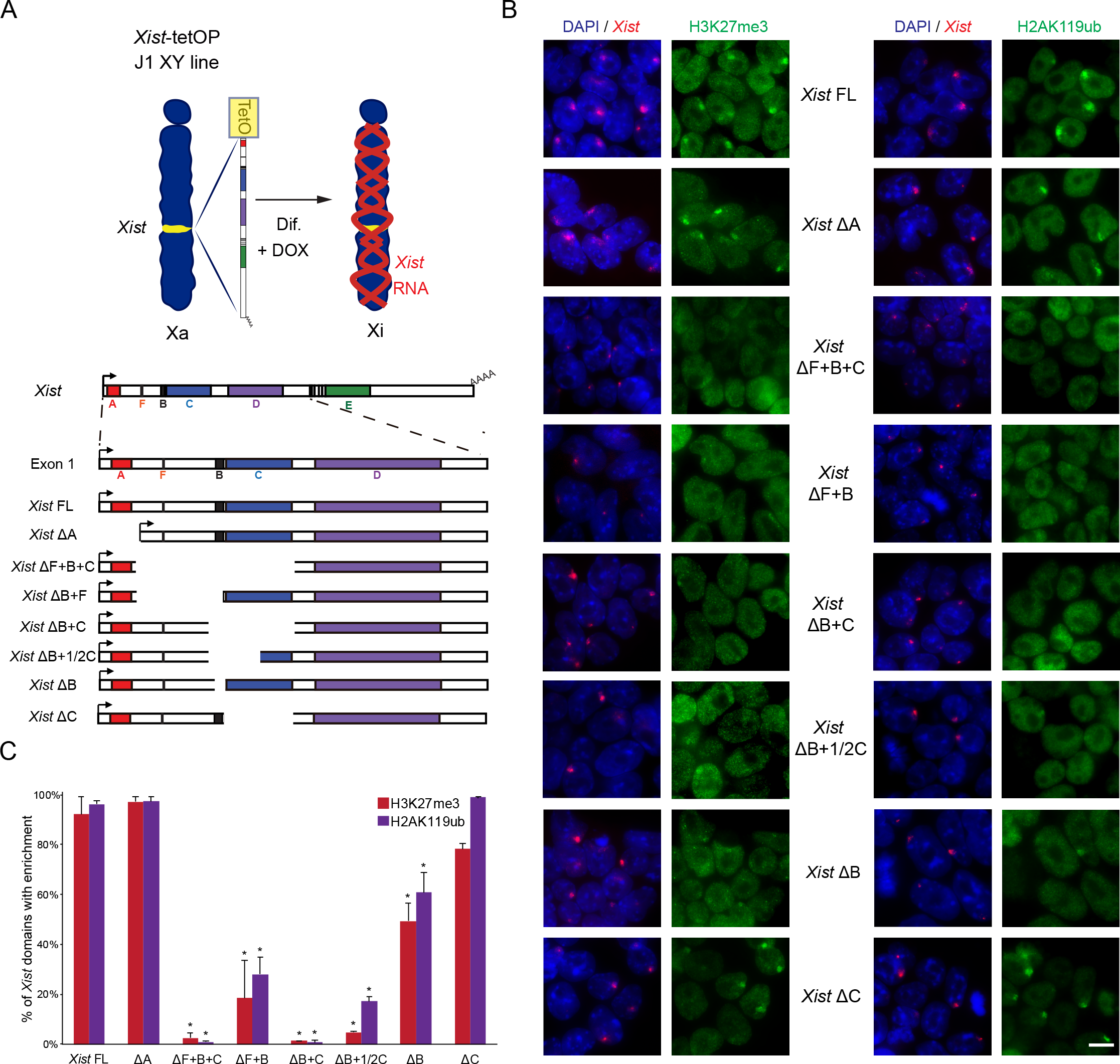
Lack of H3K27me3 and H2AK119ub enrichment over the X chromosome in the absence of *Xist* repeats B and C. A. Schematic representation of the novel *Xist*-TetOP mutants generated by CRISPR/Cas9 genome editing in J1 XY ESCs; the different repeats are highlighted in color boxes; Dif. – differentiation; DOX – doxycycline. B. Representative images of combined IF for H3K27me3 or H2AK119ub (green) with RNA FISH for *Xist* (red) in *Xist*-TetOP lines (for clone 1 of each mutant type) upon D2 in DOX conditions; Blue - DAPI staining; Scale bar: 10 μm. C. Graph represents the % of *Xist*-coated X chromosomes enriched for H3K27me3 or H2AK119ub in the different *Xist*-TetOP mutants (for clone 1 of each mutant type) from 2-to-4 independent experiments; A minimum of 50 *Xist*-coated X chromosomes were counted per experiment; Significant differences from unpaired Student’s *t*-test comparing mutants to *Xist* FL are indicated as * (p-value < 0.05).

The exact contributions of *Xist*’s repeat regions to XCI have remained unclear due to three main issues: (1) some of the deletions at the endogenous *Xist* locus can impair expression of the mutant allele and/or lead to skewed XCI towards the wild-type allele (Hoki et al., 2009; Lv et al., 2016; Senner et al., 2011); (2) deletions of the repeat elements can result in delocalization of *Xist* from the Xi territory, which indirectly affects gene silencing and chromatin changes (Ridings-Figueroa et al., 2017; Sunwoo et al., 2017; Yamada et al., 2015); (3) deletions performed in the context of autosomal cDNA inducible systems are difficult to interpret due to the reduced efficiency of *Xist*-mediated silencing of autosomal genes (Loda et al., 2017; Pintacuda et al., 2017; Tang et al., 2010).

In this study, we generate and analyse a series of *Xist* mutants created at the endogenous *Xist* gene under an inducible promoter. In particular, we explored the endogenous *Xist* RNA’s sequence requirements for recruitment of PRC1/PRC2 and re-assessed the relationship between the initiation of X-linked transcriptional silencing and PcG recruitment. Our results reveal that removal of both *Xist* B and C repeats, not just the B repeat as previously proposed in the context of autosomal transgenes, is necessary to fully abolish PRC1/PRC2 recruitment in the context of the X chromosome. Moreover, we provide evidence that X-linked transcriptional silencing can be induced in a PcG-defective *Xist* mutant, albeit slightly less efficiently.

## Results

### Generation of *Xist* RNA mutants for F, B and C repeats

To dissect the role of different functional RNA domains of *Xist*, particularly the RNA sequences enabling recruitment of PRC1 and PRC2 complexes to the X chromosome, we created a series of new inducible *Xist* mutants. For this we used a previously described system whereby *Xist* at its endogenous locus in J1 XY embryonic stem cells (ESCs) is driven by a tetracycline-inducible promoter (*Xist*-TetOP) that can be activated by doxycycline (DOX) (Figure 1A). This system recapitulates hallmarks of XCI, namely chromosome-wide *Xist* coating, X-linked gene silencing and heterochromatin formation (Wutz et al., 2002). We created 6 new mutants within *Xist* exon 1: ΔF+B+C, ΔF+B, ΔB+C, ΔB+1/2C, ΔB and ΔC by CRISPR/Cas9 genome editing using flanking pairs of guide RNAs (gRNAs) (Figure 1A; Material and Methods). At least two clones per type of mutation were created (Figure 1 - source data 1). The previously generated *Xist* ΔA mutant, which is silencing-defective (Wutz et al., 2002), but competent for PRC2 recruitment (da Rocha et al., 2014), was also used in this study for comparison (Figure 1A).

The newly generated *Xist* mutants were validated at the DNA and RNA level by PCR and RT-PCR and the exact deleted regions were mapped by Sanger sequencing (Figure 1 - figure supplement 1A-B). RNA Fluorescent In Situ Hybridization (FISH) analysis showed that all *Xist* mutants are able to form a *Xist* domain (on average, *Xist* ΔF+B+C and *Xist* ΔF+B have smaller domains) upon DOX induction, but not in non-induced (noDOX) conditions (Figure 1 - figure supplement 1C). The proportion of cells with a *Xist*-coated X chromosome varied somewhat between the mutant clones [*e.g.*, *Xist* FL: 45 ± 6 %; *Xist* ΔA: 53 ± 9 %; *Xist* ΔB+C: 60 ± 8 % at day 4 of differentiation in DOX conditions; Figure 1 - figure supplement 1C]. The two clones of each mutant type did not always have the same percentage of cells with *Xist* domains (Figure 1 - source data 1), suggesting that the differences between the lines are unlikely to be explained by *Xist* mutant type, but rather by the variable ability of cell lines to respond to DOX (Figure 1 - source data 1). Next, we employed this new series of *Xist*-TetOP mutants to assess their XCI and chromatin associated phenotypes.

### PcG complexes are recruited to the X chromosome thanks to B and C repeats of *Xist*

Previously, our results and those of others have shown that PRC1/PRC2 recruitment is impaired in the *Xist* ΔXN cDNA mutant that lacks 3.8Kb including the F, B and C repeats (Almeida et al., 2017; da Rocha et al., 2014). More recently, a 600bp region including the B repeat as well as the first 3 of the 14 motifs of the C repeat was deleted and reported to abrogate Polycomb recruitment, although this was in the context of autosomal *Xist* inducible cDNA transgenes (Pintacuda et al., 2017). To assess this in the context of the X chromosome, we evaluated whether the different *Xist*-TetOP mutants exhibited typical H3K27me3 and H2AK119ub foci over the *Xist*-coated X chromosome. For this, we performed combined immunofluorescence (IF)/*Xist* RNA FISH at day 2 of differentiation in the presence of DOX, a time-point where PcG recruitment reaches its maximum (da Rocha et al., 2014). Enrichment of H3K27me3 and H2AK119ub was seen at the *Xist*-coated X chromosome in most cells in the *Xist* FL and *Xist* ΔA cell lines (Figure 1B-C), consistent with previous reports (Almeida et al., 2017; da Rocha et al., 2014; McHugh et al., 2015). Interestingly, lack of the *Xist* C repeat alone did not significantly affect H3K27me3 or H2AK119ub enrichment. In contrast, no H3K27me3 or H2AK119ub accumulation was observed in the *Xist* ΔF+B+C mutant (Figure 1B-C), which is equivalent to the *Xist* ΔXN cDNA mutant (Almeida et al., 2017; da Rocha et al., 2014; Pintacuda et al., 2017; Wutz et al., 2002). Importantly, all *Xist*-TetOP mutants for which B repeat was absent showed a statistically significant decrease in H3K27me3 and H2AK119ub over the *Xist*-coated X chromosome (Figure 1B-C). Nevertheless, a slight enrichment of these marks was still seen in around half of *Xist* domains in *Xist* ΔB, to a lesser degree in *Xist* ΔF+B and even less in *Xist* ΔB+1/2C, which lacks 62% of the C repeat. In the *Xist* ΔB+C and ΔF+B+C mutants, no H3K27me3 and H2AK119ub enrichment was observed (Figure 1B-C). These results were confirmed in the clone 2 for each mutant type (Figure 1 - source data 2). The defects in the enrichment of PcG-associated histone modifications were associated with reduced recruitment of PRC2 (EZH2) and its co-factor (JARID2) and of PRC1 (RING1B) (Figure 1 - figure supplement 2; Figure 1 - source data 2). These defects were more pronounced, likely because histone marks are stably maintained while the PcG complexes are dynamically recruited. All in all, our results show that PRC1 and PRC2 require the same *Xist* RNA modules to enable recruitment to the X chromosome. This is consistent with the dependence of PRC2 recruitment on the non-canonical PRC1 (Almeida et al., 2017; Cooper et al., 2016; da Rocha et al., 2014; Pintacuda et al., 2017). Furthermore, we show that in the context of endogenous *Xist* locus, the deletion of both B and C repeats is needed to completely abrogate PcG recruitment. This is a significantly bigger region than that necessary to cause the same defect in the context of autosomal *Xist* transgene integrations (2.1Kb versus 0.6Kb) (Pintacuda et al., 2017). Thus, the severity of phenotypes seems to depend on the chromosomal context where *Xist*-dependent gene silencing is induced.

### *Xist* ΔB+C RNA does not interact with PCGF3/5-PRC1

To obtain mechanistic insight into why *Xist* ΔB+C RNA does not recruit PcG proteins globally, we analyzed the protein interactome of the *Xist* ΔB+C RNA using ChIRP-MS (RNA-binding proteins by mass spectrometry). Previously, ChIRP-MS identified 81 *Xist* protein partners, three of which (SPEN, WTAP and RNF20) bind to the A repeat (Chu et al., 2015). We performed ChIRP-MS on both *Xist* FL and *Xist* ΔB+C cells in induced (DOX) conditions at day 3 of differentiation as previously performed for *Xist* FL and *Xist* ΔA differentiated ES cells (Chu et al., 2015). As a negative control, *Xist* FL ES cells was also differentiated in noDOX conditions (Figure 2A). We confirmed that *Xist* RNA was retrieved after ChIRP procedure in DOX, but not in noDOX conditions (Figure 2 - figure supplement 1A). The *Xist* RNA levels recovered from *Xist* ΔB+C were higher than for *Xist* FL induced cells (Figure 2 - figure supplement 1A), as expected given the greater number of cells presenting a *Xist*-coated X chromosome in *Xist* ΔB+C (50.9%) than *Xist* FL (24.0%) as measured by RNA FISH in this experiment (Figure 2 - figure supplement 1B). Proteins retrieved by *Xist* ChIRP were separated by electrophoresis (Figure 2 - figure supplement 1C) and sent for identification by liquid chromatography-tandem mass spectrometry (LC-MS/MS).

**Figure 2.**
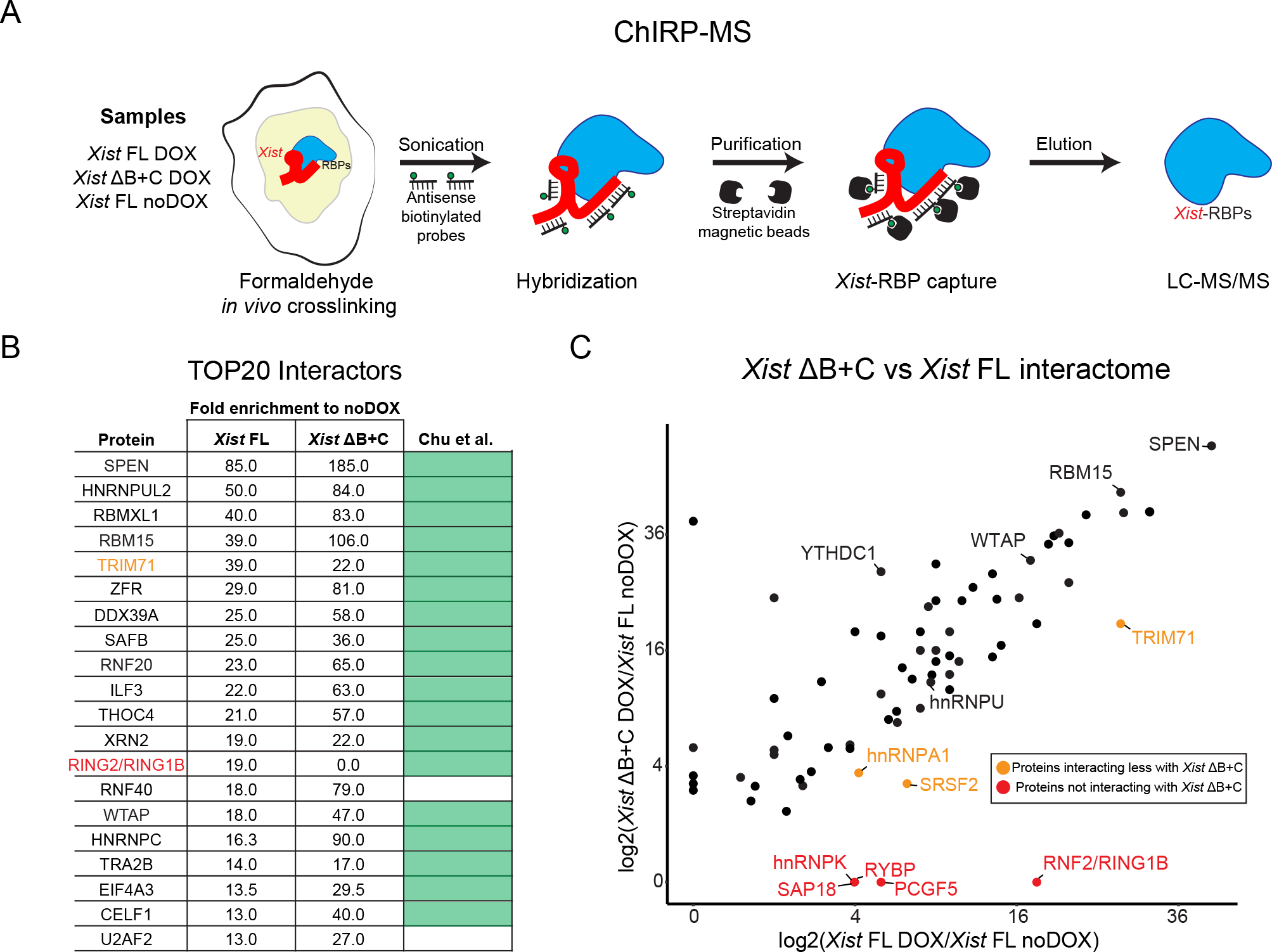
Absence of PCGF3/5-PRC1 proteins from the *Xist* ΔB+C RNA protein interactome. A. Scheme of the ChIRP-MS workflow performed on *Xist* FL (DOX and noDOX conditions) and *Xist* ΔB+C (DOX) at day 3 of differentiation; RBP - RNA binding protein. B. Top 20 protein hits from the ChIRP-MS of *Xist* FL; The ranking was based on fold-enrichment of *Xist* FL DOX versus *Xist* FL noDOX; Weakly annotated protein isoforms with an Annotation score in UniprotKB < 3 (out of 5) were excluded; Fold-enrichment for *Xist* B+C is also displayed for comparison; Light green boxes correspond to proteins previously described by Chu et al. (2015) as *Xist* interactors (Chu et al., 2015); protein in red (RING2/RING1B) represents a protein not found in the *Xist* ΔB+C interactome; Protein in light brown (TRIM71) is less enriched in *Xist* ΔB+C than in *Xist* FL. C. Scatter plot displaying the differences in peptide counts between *Xist* FL and *Xist* ΔB+C for the 74 out of 81 *Xist*-interactors from Chu et al., 2015 (Chu et al., 2015) with a minimum of fold-change of 2.5 in *Xist* FL or *Xist* ΔB+C; Shown is the log2 fold change of peptide counts of each mutant in DOX conditions compared with the *Xist* FL in noDOX conditions; proteins retrieved by both *Xist* FL and *Xist* ΔB+C ChIRPs with a proposed role in XCI such as SPEN, RBM15, WTAP, YTHDC1 and hnRNPU are indicated; light brown dots mark proteins more represented in *Xist* FL than in *Xist* ΔB+C ChIRPs, while red dots display proteins which are only retrieved by *Xist* FL ChIRP.

Previously described *Xist* protein interactors (Chu et al., 2015) were found among the top hits in *Xist* FL RNA (and also in *Xist* ΔB+C RNA) confirming the success of the ChIRP-MS experiment (Figure 2 - source data 1). Indeed, considering the 20 top hits for *Xist* FL RNA after filtering out weakly annotated protein isoforms, 18 of them are in the Chu et al. list (Chu et al., 2015). Of these top 20 hits all were shared with *Xist* ΔB+C RNA, with the notable exception of the PRC1 component RING2/RING1B (Figure 2B; Figure 2 - source data 1). Overall higher fold enrichment for the remaining factors in *Xist* ΔB+C compared to *Xist* FL is consistent with the increased yield of *Xist* RNA and proteins retrieved from mutant cells (Figure 2B; Figure 2 - figure supplement 1A & C; Figure 2 - source data 1).

By focusing on the 81 hits previously identified by Chu et al., (2015) (Chu et al., 2015), we compared the protein interactomes of *Xist* FL and *Xist* ΔB+C RNAs (Figure 2C). Consistently with our previous analysis, we detected the majority of published hits to interact with both *Xist* FL and *Xist* ΔB+C (74 out of 81 with a minimum of 2.5 DOX/noDOX fold-change in one of the samples) (Figure 2 - source data 1). SPEN and many other proteins with a proposed role in long-range gene silencing were present in the *Xist* ΔB+C interactome, such as members of the m6A RNA methyltransferase machinery (RBM15, WTAP and YTHDC1) and proteins involved in *Xist* spreading such as the hnRNPU matrix attachment protein (Figure 2C). In contrast, 5 proteins were absent from the *Xist* ΔB+C interactome (Figure 2C), including the three members of non-canonical PRC1 present in Chu et al.’s list – RNF2/RING1B, RYBP and PCGF5. We also found that PCGF3, which was not in the original Chu et al.’s list, present in the *Xist* FL interactome, but lacking from the *Xist* ΔB+C interactome in our ChIRP-MS experiment (Figure 2 - source data 1). Furthermore, hnRNPK, a RNA binding domain previously linked to *Xist*-induced PCGF3/5-PRC1 recruitment, was also not detected in *Xist* ΔB+C ChIRP-MS (with the exception of two poorly annotated isoforms) (Figure 2 - source data 1). A histone deacetylase complex subunit SAP18 was also lacking from the *Xist* ΔB+C interactome, while three other proteins were found to bind more weakly to *Xist* ΔB+C than to *Xist* FL RNA: TRIM71 (an E3 ubiquitin-protein ligase); SRSF2 (Serine and Arginine Rich Splicing Factor 2) and hnRNPA1 (Heterogeneous Nuclear Ribonucleoprotein A1) (Figure 2C; Figure 2 - source data 1).

In conclusion, *Xist* ΔB+C and *Xist* FL RNAs shares most of their protein interactome with few exceptions such as proteins of the PCGF3/5-PRC1 complex. Combined with previous results on *Xist* ΔA (Chu et al., 2015), these data illustrate the modular organization of *Xist* lncRNA, with RNA motifs interacting independently with different proteins and possibly performing distinct functions. The absence of PCGF3/5-PRC1 from the *Xist* ΔB+C interactome explains the global lack of H2AK119ub and concomitant loss of H3K27me3 enrichment over the X chromosome (Figure 1B-C), consistent with the hierarchical model proposed for PRC1/PRC2 recruitment (Almeida et al., 2017).

### Residual accumulation of PcG marks over active genes in the *Xist* ΔB+C-coated X chromosome

To assess the lack of H3K27me3 and H2AK119ub accumulation at the chromosome-wide level in the *Xist* ΔB+C mutant differentiating ES cells, we performed native ChIP-seq (nChIP-seq) for these marks. Both marks were assessed after 2 days of differentiation in DOX and noDOX conditions in biological duplicates for *Xist* ΔB+C mutant cells and compared to the results previously obtained for *Xist* FL cells (Zylicz, Bousard et al., *in press*). At autosomal sites, similar patterns of enrichment for the PcG marks were observed for all the samples in both DOX and noDOX conditions (*e.g*. *HoxC* cluster in Figure 3 - figure supplement 1A). At the level of the X chromosome, we observed a general loss of H3K27me3 and H2AK119ub accumulation in the *Xist* ΔB+C mutant in clear contrast to *Xist* FL (Figure 3A; Figure 3 - figure supplement 1B). However, we noted some residual accumulation for both marks in *Xist* ΔB+C at active, gene-dense regions (Figure 3A). We therefore evaluated enrichment at specific types of genomic regions: intergenic, promoters and gene bodies which were initially active in noDOX conditions (herein called as active promoters and active gene bodies, respectively; see Material and Methods for definition). Consistent with the chromosome-wide analysis, at intergenic windows, we observed a striking lack of H3K27me3 and H2AK119ub enrichment upon induction of *Xist* ΔB+C when compared to *Xist* FL RNA (Figure 3B). In contrast, slight enrichment of both PcG-associated marks was detected upon *Xist* ΔB+C induction at active promoters and gene bodies, in particular for H3K27me3 over active promoters (Figure 3B; Figure 3 - figure supplement 1C). This enrichment of both marks was significantly lower than that observed in *Xist* FL expressing cells (Figure 3B), as can be visualized using average plots around transcriptional start sites (TSS) of active genes (Figure 3C). We also normalized our data for the percentage of cells presenting *Xist*-coated chromosomes, based on RNA FISH analysis (Figure 3 - figure supplement 2A) and obtained similar results (Figure 3 - figure supplement 2B). Examples of typical nChIP-seq profiles are depicted in Figure 3D showing a gene with lack of accumulation for PcG marks (*Lamp2*), and a second gene (*Rlim/Rnf12*) with clear H3K27me3/H2AK119ub enrichment around the promoter in the induced *Xist* ΔB+C mutant cells. In conclusion, we observed no enrichment of H3K27me3 and H2AK119ub at intergenic regions upon expression of *Xist* ΔB+C RNA, but a mild accumulation is seen over some active promoters and to a lesser extent at gene bodies. As genes represent only a small fraction of the X chromosome, this is probably why we could not detect their enrichment in the IF/RNA FISH experiments. The reasons behind this mild enrichment of H3K27me3 and H2AK119ub at some X-linked genes in the *Xist* ΔB+C mutant that cannot bind PCGF3/5-PRC1 proteins are unclear, but could be due to the transcriptional silencing of these genes.

**Figure 3.**
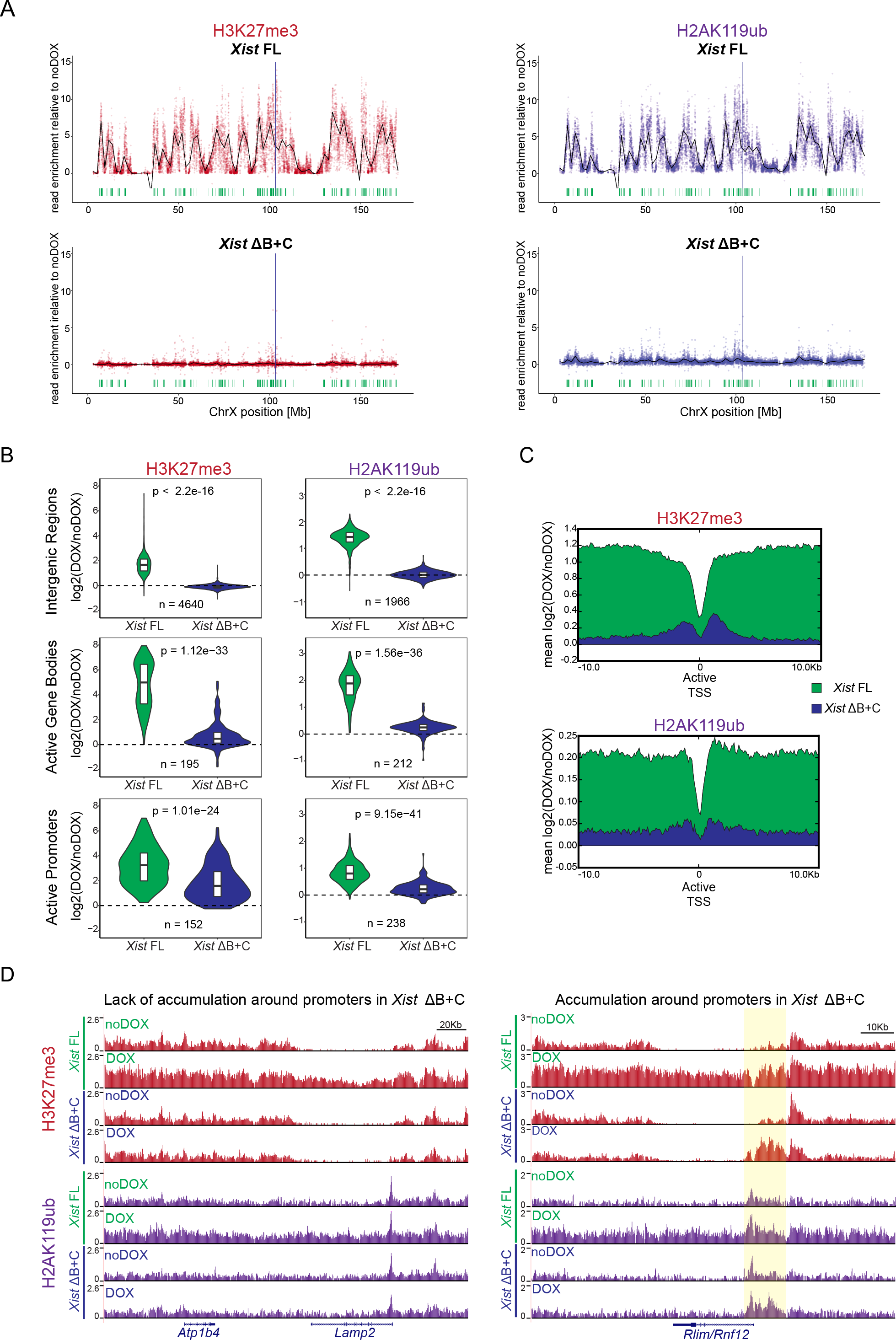
nChIP-seq reveals chromosome-wide absence of H3K27me3 and H2AK119ub enrichment, but residual enrichment at active genes in the *Xist* ΔB+C. A. Plots showing H3K27me3 and H2AK119ub accumulation over the X chromosome in *Xist* FL and *Xist* ΔB+C with upon DOX induction at day 2 (D2) of differentiation; Each dot represents a single 10Kb window and its enrichment relative to noDOX condition; Black line is a loess regression on all windows; *Xist* locus is represented by a blue long line, active genes by green lines. B. Violin plots quantifying H3K27me3 and H2AK119ub enrichment over intergenic regions, active promoters and active gene bodies in the X chromosome in *Xist* FL and ΔB+C cell lines at D2 upon DOX induction; Shown is the log2 fold change of DOX vs noDOX conditions; n = indicates the number of regions/genes analyzed; p-values were calculated using paired Wilcoxon test, comparing *Xist* FL and *Xist* ΔB+C cell lines. C. Average plots showing the mean enrichment of H3K27me3 (top) and H2AK119ub (bottom) over all X-linked active transcriptional start sites (TSS); Shown is the mean of normalized log2 enrichment of DOX vs noDOX in both *Xist* FL and *Xist* ΔB+C cell lines. D. Genome browser plots showing H3K27me3 (top) and H2AK119ub (bottom) enrichments in a region encompassing the inactive *Atp1b4* and the initially active *Lamp2* genes within the XqA3.3 region and the initially active *Rlim/Rnf12* gene at the XqD region; Region around the promoter of *Rlim/Rnf12* is highlighted in yellow.

### *Xist* ΔB+C RNA is able to initiate long-range transcription silencing along the X chromosome

To assess the degree to which transcriptional silencing could be induced in the *Xist* ΔB+C expressing cells, we evaluate expression from X-linked genes. We initially performed nascent transcript RNA FISH combined with *Xist* RNA FISH for X-linked genes (*Pgk1* and *Lamp2* at D2; *Pgk1* and *Rlim*/*Rnf12* at D4) in different *Xist* mutant lines (Figure 4A-B; Figure 4 - source data 1). As expected, *Xist* ΔA RNA was entirely defective in silencing for the assessed X-linked genes. In striking contrast, all the other *Xist* mutants (ΔF+B+C, ΔF+B, ΔB+C, ΔB+1/2C, ΔB & ΔC) were able to silence these genes at levels approximately similar to *Xist* FL RNA (Figure 4A-B; Figure 4 - source data 1). Similar results were obtained for the second clone of each mutant (Figure 4 - source data 1). Corroborating the nascent-transcript RNA FISH data, we also noted significant reduction in cell survival upon prolonged DOX induction (≥ 5 days) for *Xist* FL and all the mutants with the exception of *Xist* ΔA RNA (data not shown). This is consistent with efficient XCI in XY ESCs, resulting in functional nullisomy for the X chromosome, and thus cell death. Interestingly, a mild relaxation of silencing could be seen for some genes, as for example, the *Lamp2* gene at D2 in PcG-defective *Xist* ΔF+B+C and *Xist* ΔB+C, but not in *Xist* FL or *Xist* ΔB and *Xist* ΔC (Figure 4A).

**Figure 4.**
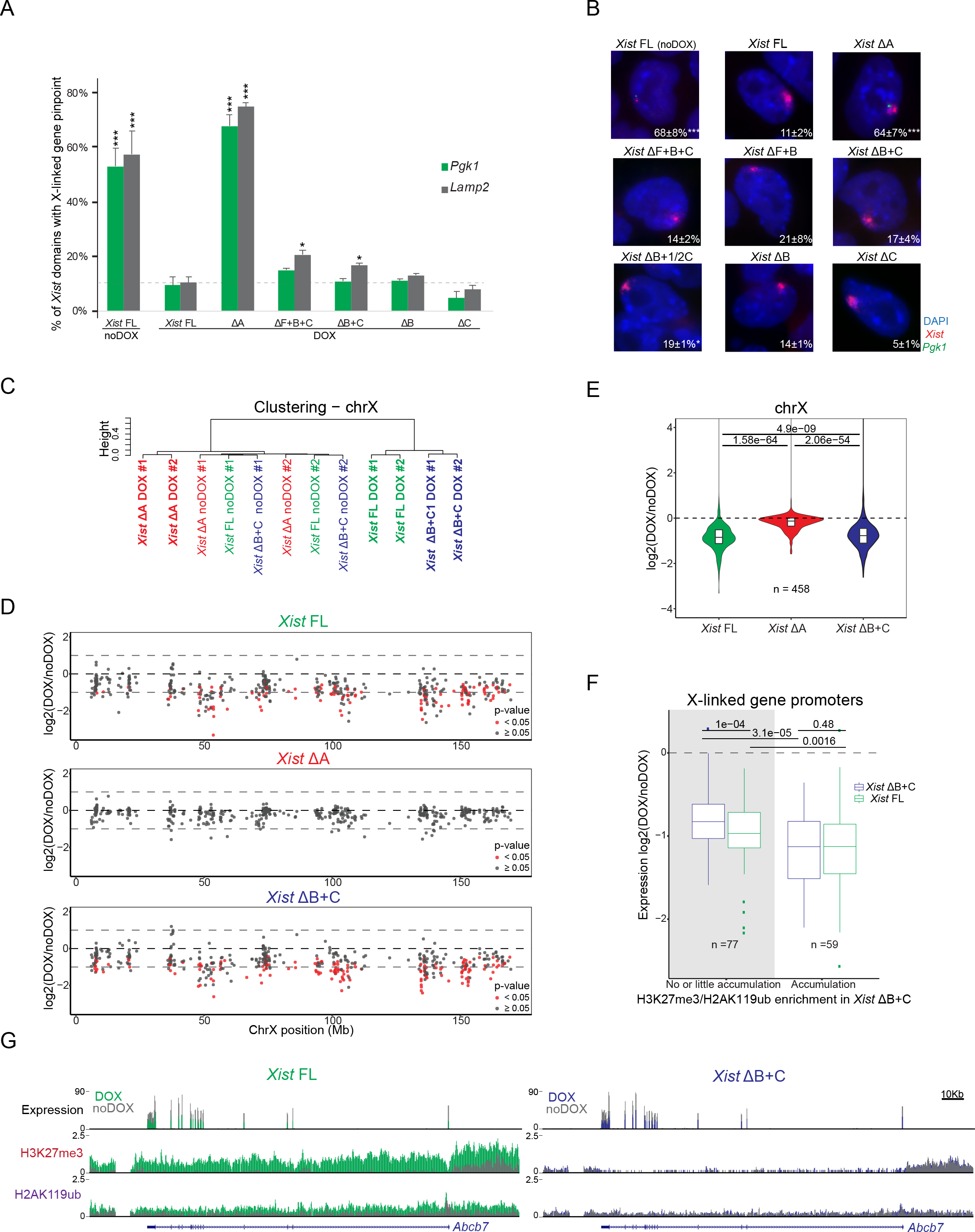
*Xist* ΔB+C is able to initiate X chromosome-wide transcriptional silencing with no or residual Polycomb recruitment. A. Graph represents the mean % + S.E.M. of *Xist*-coated chromosomes presenting an active *Pgk1* or *Lamp2* gene as determined by RNA FISH (as represented in B) at D2 in the presence of DOX (*Xist* FL was also used in noDOX conditions) in the different *Xist*-TetOP mutants; each bar represents the mean from to 2-to-4 independent experiments; A minimum of 50 *Xist*-coated chromosomes were counted per experiment; For *Xist* FL noDOX a minimum of 100 cells (which do not have *Xist*-coated chromosome) were counted; Significant differences compared with *Xist* FL (DOX) are indicated as * (p-value < 0.05) or *** (p-value < 0.01), unpaired Student’s *t*-test; dashed line marks the mean percentage of silencing for the *Lamp2* gene in *Xist* FL DOX. B. Representative RNA FISH images for *Xist* (red) and nascent-transcript of *Pgk1* (green) in *Xist*-TetOP lines at day 4 of differentiation in the presence of DOX (*Xist* FL is also shown in noDOX conditions); DNA stained in blue by DAPI; Numbers represent % of *Xist*-coated X-chromosomes ± S.E.M. with active *Pgk1* gene (except for *Xist* FL noDOX, where numbers represent % of cells with *Pgk1* active gene); The values represent 2-to-4 independent experiments, where a minimum of 50 *Xist*-coated chromosomes were counted per experiment; Significant differences compared with *Xist* FL (DOX) are indicated as * (p-value < 0.05) or *** (p-value < 0.01), unpaired Student’s *t*-test. C. Clustering analysis of the normalized RNA-seq counts on the X-chromosome (chrX) for all the duplicates of *Xist* FL, *Xist* ΔA and *Xist* ΔB+C in DOX and noDOX conditions. D. Plots displays the log2(fold-change) in the expression of X-linked genes along the chrX comparing DOX versus noDOX samples for *Xist* FL, *Xist* ΔA and *Xist* ΔB+C at day 2 of differentiation; red dots correspond to genes which are differently expressed in DOX vs noDOX (p < 0.05, Limma *t*-test), while black dots represent genes which are not differentially expressed between the two conditions (p ≥ 0.05).
E. Violin plots displaying the average log2(fold-change) in gene expression between DOX and noDOX conditions on the chrX in *Xist* FL, *Xist* ΔA and *Xist* ΔB+C at day 2 of differentiation; p-values were calculated using paired Wilcoxon test; n = indicates the number of genes analyzed. F. Box plots displaying the log2(DOX/noDOX) fold-change in expression of X-linked genes in *Xist* FL and *Xist* ΔB+C categorized according to the enrichment of H3K27me3 and H2AK119ub marks at promoters in *Xist* ΔB+C upon DOX induction (with no or little accumulation vs accumulation); p-values between samples were calculated using paired Wilcoxon test; n = indicates the number of genes analyzed. G. Genome browser plots showing RNA-seq reads, H3K27me3 and H2AK119ub nChIP reads around the *Abcb7* gene for *Xist* FL (left) and *Xist* ΔB+C (right) at day 2 of differentiation in both DOX and noDOX conditions.

To assess the full extent of transcriptional silencing of X-linked genes in the absence of *Xist*-mediated PcG recruitment, we examined RNA-seq on biological duplicates of *Xist* FL, *Xist* ΔA (silencing-defective) and *Xist* ΔB+C (PcG-defective) in DOX and noDOX conditions at day 2 of differentiation. We confirmed robust *Xist* upregulation upon DOX treatment and found no reads mapping to the deleted regions in both mutant lines (Figure 4 - figure supplement 1A). First, we evaluated whether the percentage of total X-chromosome specific RNA-seq reads changed before and upon induction. While no changes were observed for the silencing-defective *Xist* ΔA cell line, the percentage of X-chromosome specific reads decreased in both *Xist* FL and *Xist* ΔB+C cell lines upon DOX induction (Figure 4 - figure supplement 1B). Clustering analysis based on X-linked gene expression shows that DOX-induced samples of *Xist* FL and *Xist* ΔB+C segregate from the noDOX samples and DOX-induced *Xist* ΔA samples (Figure 4C). Furthermore, both *Xist* FL and *Xist* ΔB+C RNAs, but not *Xist* ΔA, were able to silence most genes throughout the X chromosome (Figure 4D). This is consistent with our previous nascent transcript RNA FISH analysis (Figure 4A-B). When we compared the average degree of silencing, we observed a slight relaxation of X-linked gene silencing in the *Xist* ΔB+C mutant when compared to *Xist* FL expressing cells (Figure 4E). This becomes more evident when data are adjusted for the percentage of cells presenting *Xist*-coated chromosomes as judged by RNA FISH (Figure 4 - figure supplement 1C-D). Nonetheless, this effect on gene silencing was significantly milder than that observed for *Xist* ΔA mutant expressing cells (Figure 4E and Figure 4 - figure supplement 1D). The slight silencing defect in *Xist* ΔB+C expressing cells appears to be a chromosome-wide effect, since we could not pinpoint specific genes driving the differences in silencing efficiency between *Xist* ΔB+C and *Xist* FL (Figure 4 - figure supplement 1E). All in all, these results show that *Xist* ΔB+C RNA is able to silence X-linked genes but the degree of overall silencing is less effectively initiated and/or maintained.

Finally, we wished to explore the relationship between PcG recruitment at promoters and initiation of X-linked gene silencing, given the slight enrichment of PcG marks over some X-linked genes in *Xist* ΔB+C induced cells (Fig. 3B-C). To address this, we categorized X-linked genes by their degree of silencing based on expression fold-change differences between DOX and noDOX conditions for both *Xist* FL and *Xist* ΔB+C. In both cases, accumulation of PcG marks at promoters correlated with the level of gene silencing (Figure 4 - figure supplement 1F). Within each of these categories of similarly silenced genes, H3K27me3 and H2AK119ub enrichment were significantly lower in *Xist* ΔB+C when compared to *Xist* FL (Figure 4 – figure supplement 1F). We next assessed whether genes that do not accumulate PcG marks upon *Xist* ΔB+C induction were silenced. We found 77 X-linked genes that accumulate little or no H3K27me3 and H2K119Aub marks at their promoters specifically in *Xist* ΔB+C induced cells (Figure 4 – figure supplement 1G). These genes were nevertheless significant silenced upon induction of the *Xist* ΔB+C RNA (Figure 4F) as exemplified by the *Abcb7* gene (Fig. 4G). This suggests that PcG recruitment seems to be dispensable for initiating silencing of these genes. We noted, however, a slight silencing relaxation of these 77 genes when compared to *Xist* FL. Also, on average, these genes silenced less well than genes accumulating PcG marks in the mutant and *Xist* FL (Figure 4). This implies that either PcG recruitment is needed to stabilize silencing initially imposed by other factors or that its mild, local enrichment of H3K27me3 and H2AK119ub is simply a consequence of X-linked gene silencing in *Xist* ΔB+C. The passive recruitment model is consistent with the fact that gene promoters accumulating PcG marks in *Xist* ΔB+C (and *Xist* FL) are enriched for CpG content (Figure 4 -figure supplement 1G-H). This feature is thought to promote PcG deposition at silenced promoters (Davidovich et al., 2013; Mendenhall et al., 2010; Riising et al., 2014). In conclusion, we believe our data points to a model whereby *Xist*-mediated PcG accumulation via the B+C repeat region is not the initial driving force causing X-linked transcriptional silencing for most genes (Figure 5).

**Figure 5.**
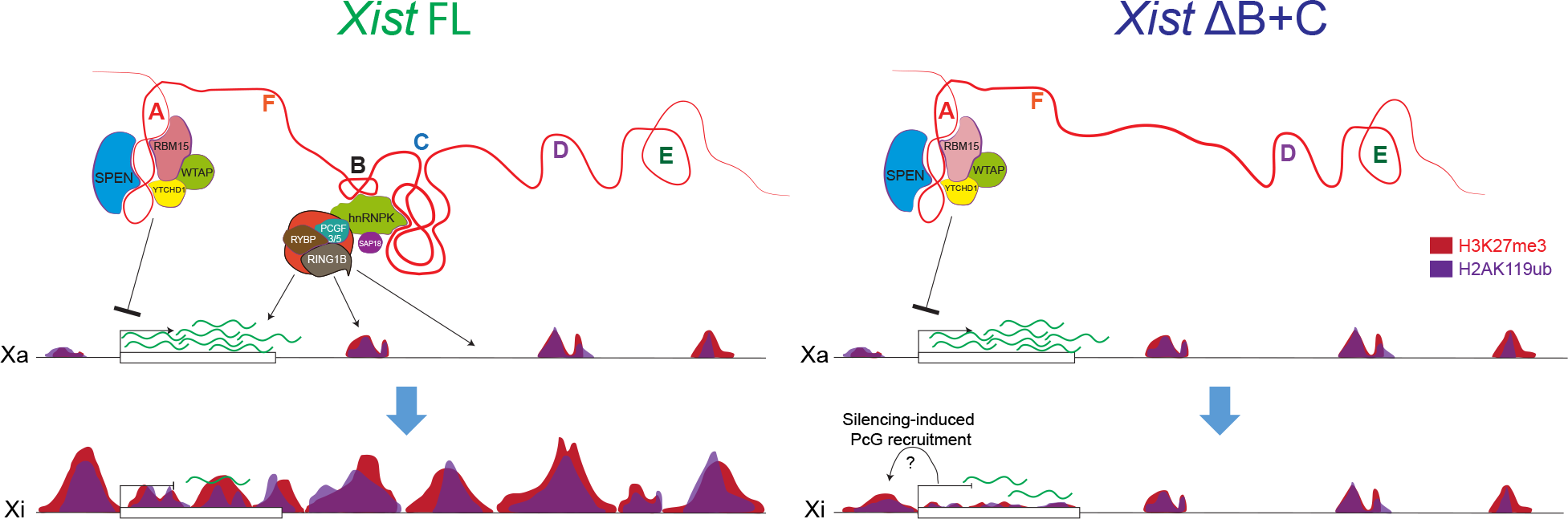
Working model for *Xist*-mediated PcG recruitment influence on transcriptional silencing based on the phenotypes of the *Xist* ΔB+C mutant. SPEN and proteins of the m6A RNA methylation machinery interact with the A repeat to initiate X-linked gene silencing; PCGF3/5-PRC1 recruitment via hnRNPK interaction with the B and C repeats is responsible for the accumulation of H2AK119ub and concomitant enrichment of the PRC2-mark H3K27me3 over the entire X-chromosome. In the absence of B and C repeats, there is no enrichment of PcG marks in intergenic regions, but a slight increase at silencing X-linked genes is seen; this could be caused by passive recruitment induced by gene silencing; nevertheless, recruitment of these marks are necessary to stabilize the initial silencing mediated by the A repeat interactors.

## Discussion

We found that chromosome-wide transcriptional silencing and PRC1/PRC2 recruitment rely primarily, respectively, on the A repeat and B+C *Xist* RNA repeats, on the X chromosome undergoing inactivation. Our analysis indicates that initiation of X-linked gene silencing can occur without *Xist*-induced chromosome-wide PRC1/PRC2 recruitment. However, PRC1/PRC2 seems to be necessary to stabilize the repressive state of some genes.

The inducible *Xist* mutants we have generated in this study represent a useful model for the study of individual *Xist* modules in the initiation of XCI, with *Xist* induction occurring at its endogenous location rather than at autosomal locations (Pintacuda et al., 2017; Loda et al., 2017; Tang et al., 2010). These *Xist* mutants allowed us to study the function of F, B and C repeats, previously reported to be important for PRC1/PRC2 recruitment (Almeida et al., 2017; da Rocha et al., 2014). We show here that a *Xist* ΔC mutant has no obvious defect in *Xist* RNA coating of the X chromosome and PcG recruitment. This contrasts with previous findings suggesting a role for the C repeat in *Xist* localization (Sarma et al., 2010) although this could be due to differences in the cell type examined (somatic cells), and the technology used (locked nucleic acids - LNAs) to destabilize the C repeat. We also show that our *Xist* mutants lacking the B repeat have impaired PRC1 and PRC2 recruitment to the X chromosome, consistent with the recent finding implicating a 600 bp region containing the B repeat (and 3 out of 14 motifs of the C repeat) on PRC1/PRC2 recruitment in an autosomal context (Pintacuda et al., 2017). However, we found that lack of B repeat alone is unable to fully compromise PRC1/PRC2 recruitment on the X chromosome. In our mutants, complete absence of PRC1/PRC2 recruitment as judged by IF/RNA FISH, is seen only if both B and C repeats are deleted. Interestingly, B and C repeats correspond precisely to the binding sites of the RNA binding protein hnRNPK as mapped by iCLIP: B repeat represents the stronger binding region for hnRNPK within *Xist*, but this protein also interacts all along the C repeat (Cirillo et al., 2016). hnRNPK was recently proposed to be an important player in mediating *Xist-*dependent recruitment of PCGF3/5-PRC1 and PRC2 to the X chromosome (Chu et al., 2015; Pintacuda et al., 2017). In accordance with this, we found that hnRNPK, alongside with non-canonical PRC1 members, is lost from the *Xist* ΔB+C protein interactome as revealed by ChIRP-MS.

The *Xist* ΔB+C mutant cannot bind PCGF3/5-PRC1 and is able to cause chromosome-wide transcriptional silencing in contrast to the silencing-defective *Xist* ΔA mutant. This can be explained, at least in part, by the interaction of *Xist* ΔB+C RNA with factors involved in X-linked gene silencing, such as SPEN, RBM15 and WTAP (Chu et al., 2015; McHugh et al., 2015; Patil et al., 2016). However, a slight relaxation of transcription silencing is still seen for the *Xist* ΔB+C mutant RNA. It is a weaker phenotype than the previously reported decrease of transcriptional silencing seen by PcG-defective *Xist* mutant transgenes on autosomes (Pintacuda et al., 2017). However, the overall decrease in *Xist*-mediated gene silencing efficiency in an autosomal context (Loda et al., 2017; Tang et al., 2010) might render autosomal genes more susceptible to PcG loss. In any case, given the slight relaxation of transcriptional silencing in *Xist* ΔB+C, this data suggests that *Xist*-dependent global PcG recruitment to the X chromosome, which is not directed specifically to genes, will be important to guarantee a stable inactive state.

Our nChIP-seq data confirmed the lack of global enrichment of PcG marks of the X-chromosome, but revealed a mild enrichment of these marks around the promoters and gene bodies of some X-linked genes in *Xist* ΔB+C expressing cells. This suggests that PcG marks may be laid down on the X chromosome in more than one way. One possibility is that another region of *Xist* mediates PcG recruitment to these genes. Although the A repeat has been previously implicated in PRC2 recruitment (Maenner et al., 2010; Zhao et al., 2008), the specificity of such an interaction is unclear (Brockdorff, 2013; Davidovich et al., 2015). Furthermore, the PRC2 core components have not been identified to bind to *Xist* RNA in different proteomics searches of the *Xist* interactome (Chu et al., 2015; McHugh et al., 2015; Minajigi et al., 2015). Another possibility is that low levels of PcG proteins may simply be recruited to promoters and gene bodies as a consequence of gene silencing following *Xist* RNA coating. Taking advantage of comparable nChIP-seq and RNA-seq data sets in *Xist* FL and *Xist* ΔB+C, we detected multiple genes which were silenced and yet with no or residual accumulation of H3K27m3 or H2AK119ub at their promoter regions in *Xist* ΔB+C induced cells. This suggests that *Xist*-mediated gene silencing can occur in the absence of PcG recruitment, at least, for a subset of X-linked genes. The recruitment of PcG at X-linked genes could be secondary to transcriptional silencing in PCGF3/5-PRC1-unbound *Xist* ΔB+C. It has been proposed that PcG recruitment to active promoters and gene bodies could be passive upon their silencing, in the sense that the PcG system will operate on any transcriptionally inactive, GC-rich locus (Davidovich et al., 2013; Mendenhall et al., 2010; Riising et al., 2014). Interestingly, our results with *Xist* ΔB+C also clearly indicate that transcriptional silencing is not sufficient to recruit PcG on X-linked genes to the same extent as *Xist* FL. Thus, lncRNA-directed PcG recruitment, which mechanistically might differ from the passive recruitment to silencing genes, is necessary for proper PcG targeting in the context of XCI.

In conclusion, our results reinforce the idea that *Xist* is a multi-tasking RNA molecule with several structural and regulatory modules (Lu et al., 2016) that have different functions. We also show that initiation of *Xist*-mediated transcriptional silencing can occur in the absence of *Xist*-mediated PcG recruitment. Our work places *Xist*-mediated PcG recruitment as an important player during XCI needed to sustain initiation of PcG-independent gene silencing.

## Materials and Methods

### ES cell lines

The previously published *Xist*-tetOP (herein *Xist* FL) and *Xist*ΔSX-tetOP (herein *Xist* ΔA) XY ES cells (Wutz et al., 2002) were adapted and maintained in feeders-free classic ES cell medium - DMEM media containing 15% fetal bovine serum (FBS), 10^3^U/mil leukaemia inhibitor factor (LIF), 10^−4^ mM 2-mercaptoethanol, 50U/ml penicillin and 50μg/ml of streptomycin (Gibco). *Xist* FL ES cell line was used to generate several *Xist* mutants: ΔF+B+C, ΔF+B, ΔB+C, ΔB+1/2C, ΔB and ΔC (see Generation of *Xist*-TetOP mutants by CRISPR-Cas9 genome editing).

All ES cells were grown at 37°C in 8% CO_2_ and medium was changed daily. Inducible expression of *Xist* driven by a TetO promoter was achieved by adding DOX (1.5μg/ml) while differentiating the ES cells in LIF withdrawal medium - DMEM media containing 10% FBS, 10^−4^mM 2-mercaptoethanol and 50U/ml penicillin and 50μg/ml of streptomycin (Gibco), for 2 to 5 days, depending on the experiment.

### Generation of *Xist*-TetOP mutants by CRISPR/Cas9 genome editing

To generate *Xist*-TetOP mutants, 4×10^5^ cells were co-electroporated with 2.5μg each of two pX459 plasmids (Addgene) expressing the Cas9 endonuclease and chimeric guide RNAs (gRNAs) flanking the region to delete using a Neon Transfection System (Thermo Fisher Scientific). The sequences of the gRNAs used to generate each type of *Xist* mutant are shown in Supplementary file 1. To pick single ES cell clones containing the desired mutations, ES cells were separated by limiting dilution. As soon as visible, single colonies were picked under a microscope and screened for deletion by PCR and absence of the wild-type band with the primers depicted in Supplementary file 1. Positive clones of each *Xist* mutant type were expanded and further validated for the mutation and absence of the wild-type band. Amplicons from the deletion PCR were gel-purified using NZYGelpure kit (NZYTech) and sequenced by the Sanger method (GATC -Eurofins Genomics) (Figure 1 – figure supplement 1A; Figure 1 - source data 1) using either the forward or the reverse primers (Supplementary file 1). PCR across the deleted regions within *Xist* exon 1 were also performed and confirmed in cDNA obtained upon 4 days of differentiation in DOX conditions, while no band (or very faint bands) were obtained in noDOX conditions (Figure 1 – figure supplement 1A). *Xist*-TetOP mutants were also analyzed for expression in DOX and noDOX conditions, using primers across exon 1-to-3, upstream of the B repeat and downstream of the C repeat using the primers in Supplementary file 2, and presence or absence of the expected band was in accordance with the respective mutant analyzed (Figure 1 – figure supplement 1B).

### RT-PCR analysis

Total RNA was isolated from the different *Xist*-TetOP mutant ES cells at D4 of differentiation (from both DOX and noDOX conditions) using NYZol (NZYtech) and then DNAse I treated (Roche) to remove contaminating DNA. RNA template was reverse transcribed using the Transcriptor High Fidelity cDNA Synthesis Kit (Roche), according to the manufacturer’s instructions. cDNA was subjected to RT-PCR using the deletion primers in Supplementary file 1 for the respective *Xist*-TetOP mutants and for the analyses in Figure 1 – figure supplement 1B using the primers in Supplementary file 2 for all the *Xist*-TetOP mutants.

### RNA FISH

RNA FISH probes for *Xist* (a 19Kb genomic λ clone 510 (Chaumeil et al., 2008), *Pgk1* (a 15-16Kb genomic sequence starting 1.6Kb upstream of *Pgk1* gene up to its intron 6) (kind gift from T. Nesterova, Univ. of Oxford) (Moindrot et al., 2015), *Lamp2* (RP24-173A8 bacterial artificial chromosome – BAC) and *Rlim/Rnf12* (RP24-240J16 BAC - BACPAC Resources Center) were prepared using the Nick Translation Kit (Abbot) with red and/or green dUTPs (Enzo Life Sciences). RNA FISH was done accordingly to established protocols (Chaumeil et al., 2008) in *Xist*-TetOP mutant differentiating ES cells with minor modifications. Briefly, cells were dissociated with trypsin (Gibco) and adsorbed onto poly-l-lysine (SIGMA)-coated 22×22mm coverslips for 5 minutes (min). Cells were then fixed in 3% paraformaldehyde (PFA) in phosphate-buffered saline (PBS) for 10min at room temperature (RT) and permeabilized with 0.5% Triton X-100 diluted in PBS with 2mM Vanadyl-ribonucleoside complex (VRC; New England Biolabs) for 5min on ice. Coverslips were then washed twice in ethanol (EtOH) 70% for 5min and then dehydrated through ethanol series (80%, 95% and 100%) and air-dried quickly before hybridization with the fluorescent labelled probes. Probes were ethanol precipitated with sonicated salmon sperm DNA (and mouse *Cot1* DNA for *Lamp2* and *Rlim/Rnf12* probes), denatured at 75°C for 7min (in the case of *Lamp2* and *Rlim/Rnf12* BAC probes, they were let incubating at 37°C for 30min after denaturation to allow *Cot1* DNA to bind to the repetitive DNA present in these BACs to prevent nonspecific hybridization). *Xist* (in red or green) and one X-linked gene (*Pgk1*, *Lamp2* or *Rlim/Rnf12*) (in green when *Xist* probe was red; in red when *Xist* probe was green) probes were co-hybridized in FISH hybridization solution (50% formamide, 20% dextran sulfate, 2x SSC, 1μg/μl BSA, 10mM Vanadyl-ribonucleoside) overnight. Washes were carried out with 50% formamide/2x saline-sodium citrate (SSC), three times for 7min at 42°C and then with only 2x SCC, three times for 5min at 42°C. After the RNA FISH procedure, nuclei were stained with 4′,6-diamidino-2-phenylindole (DAPI; Sigma-Aldrich), diluted 1:5000 in 2x SCC for 5min at RT and mounted with Vectashield Mounting Medium (Vectorlabs). Cells were observed with the widefield fluorescence microscope Zeiss Axio Observer (Carl Zeiss MicroImaging) with 63x oil objective using the filter sets FS43HE, FS38HE and FS49. Digital images were analyzed with the FIJI platform (Schindelin et al., 2012). To determine the number of cells with an *Xist-*coated X chromosome, a minimum of 250 cells were counted per single experiment. To determine the expression of the different X-linked genes studied (*Pgk1*, *Lamp2* and *Rlim/Rnf12*), at least 50 cells with a *Xist*-coated X chromosome were counted in DOX conditions and, at least, 100 cells were counted in noDOX conditions per experiment.

### IF/RNA FISH

IF/RNA FISH experiments were performed as previously (da Rocha et al., 2014). *Xist* FL and mutant ES cells were differentiated for 48 hours in the presence of DOX (1.5μg/mL) on gelatin-coated 22×22mm coverslips. Cells were fixed in 3% PFA in PBS for 10min at RT, followed by permeabilization in PBS containing 0.5% Triton X-100 and VRC (New England Biolabs) on ice for 5min. After three rapid washes in PBS, samples were blocked for, at least, 15min with 5% gelatin from cold water fish skin (Sigma) in PBS. Coverslips were incubated with the following primary antibodies diluted in blocking solution at the desired concentration (H3K27me3 – Active Motif #39155 1:200; H2AK119ub – Cell Signaling #8240 1:200; JARID2 – Abcam #ab48137 1:500; RING1B - Cell Signaling #5694 1:100; EZH2 – Leica Microsystems #NCL-L-EZH2 1:200) in the presence of a Ribonuclease Inhibitor (0.8μl/mL; Euromedex) for 45min at RT (in the case of RING1B antibody, incubation lasted for 4 hours). After three washes with PBS for 5min, the coverslips were incubated with a secondary antibody (goat anti-mouse or anti-rabbit antibodies conjugated with Alexa green, red or Cy5 fluorophores diluted 1:500) for 45min in blocking solution supplemented with Ribonuclease Inhibitor (0.8μl/mL; Euromedex). Coverslips were then washed three times with PBS for 5min at RT. Afterwards, cells were post-fixed with 3% PFA in PBS for 10min at RT and rinsed three times in PBS and twice in 2x SSC. Excess of 2x SSC was removed and cells were hybridized with a *Xist* p510 probe labelled with Alexa green or red dUTPs (prepared and hybridized as mentioned in the RNA FISH protocol). After the RNA FISH procedure, nuclei were stained with DAPI (Sigma-Aldrich), diluted 1:5000 in 2x SCC for 5min at RT and mounted with Vectashield Mounting Medium (Vectorlabs). Cells were observed with the widefield fluorescence microscope Zeiss Axio Observer (Carl Zeiss MicroImaging) with 63x oil objective using the filter sets FS43HE, FS38HE and FS49. Digital images were analyzed with the FIJI platform (Schindelin et al., 2012). Enrichment of the different histone marks or PcG fluorescent signals over *Xist* cloud marked by RNA FISH were counted from at least 50 cells per single experiment.

### *Xist* ChIRP-MS

*Xist* FL (both in DOX and noDOX conditions) and *Xist* ΔB+C (DOX) cells were differentiated for 3 days. A fraction of these cells were always used to quantify levels of *Xist* induction by RNA FISH. *Xist* ChIRP-MS were conducted using a previously published protocol (Chu et al., 2015) with the following modifications: (1) around 500 million cells per ChIRP-MS experiment were collected (roughly 10-15 15cm^2^ dishes) cross-linked in 3% formaldehyde for 30min, followed by 0.125M glycine quenching for 5min; (2) All 100mg of cell pellets were then dissolved in 1ml of nuclear lysis buffer (50 mM Tris–Cl pH 7.0, 10 mM EDTA, 1 % SDS) and 880μl were sonicated in a Covaris ultrasonicator for 1 hour (20min, three times). Clarified lysates were pooled for each sample; (3) instead of RNase treatment, noDOX condition was used as control. 6μl of Riboblock RNase inhibitor was added per ml of clear lysate into the experiment and control tubes were incubated at 37°C for 30min prior to hybridization step. Final protein samples were size-separated in bis-tris SDS-PAGE gel for LC/MS-MS. Correct retrieval of *Xist* RNA after ChIRP from *Xist* FL and *Xist* ΔB+C was analyzed by RT-qPCR using three pairs of primers along *Xist* RNA [Pair 1 - Forward 1 (Fw1): GCCT CTGA TTTA GCCA GCAC, Reverse 1 (Rv1): GCAA CCCA GCAA TAGT CAT; Pair2 - Fw2: GACA ACAA TGGG AGCT GGTT, Rv2: GGAT CCTG CACT GGAT GAGT; Pair 3 - Fw3: GCCA TCCT CCCT ACCT CAGAA; Rv3: CCTG ACAT TGTT TTCC CCCT AA) and *Gapdh* as housekeeping gene (Fw: AAGG TCAT CCCA GAGC TGAA; Rv: CTGC TTCA CCAC CTTC TTGA)]. For details on ChIRP probe design, please see Extended Experimental Procedure on the previously published protocol (Chu et al., 2015).

*Xist* hits from CHIRP-MS were ranked according to *Xist* FL DOX/*Xist* FL noDOX fold-change in peptide counts. To calculate this and *Xist* ΔB+C/*Xist* FL noDOX ratios, when peptide counts for *Xist* FL noDOX samples was 0, it was considered 1 (Figure 2 - source data 1). For comparison with Chu et al. 2015 list (Chu et al., 2015), only annotated protein isoforms with an Annotation score in UniportKB ≥ 3 (out of 5) were considered with a minimum of 2.5 DOX/noDOX fold-change in one of the samples. Proteins present in the Chu’s list with DOX/no DOX ratio inferior in *Xist* ΔB+C than *Xist* FL were considered underrepresented in *Xist* ΔB+C protein interactome. Proteins with no peptide counts for *Xist* ΔB+C or with equal peptide counts to *Xist* FL noDOX, which had a DOX/noDOX ratio ≥ 4 for *Xist* FL were considered not to be part of the *Xist* ΔB+C protein interactome.

### nChIP-seq

nChIP-seq was performed in duplicates for *Xist* ΔB+C ES cells at day 2 of differentiation upon DOX and noDOX conditions and compared to results previously obtained for *Xist* FL (Zylicz, Bousard et al., *in press*). The protocol was followed as described in Zylicz, Bousard et al., *in press*. Briefly, around 3.5 million cells were used per immunoprecipitation (IP) experiment. A fraction of these cells was always used to quantify levels of *Xist* induction by RNA FISH. Ten million cells were resuspended and lysed in 90μl of Lysis Buffer (50mM Tris-HCl, pH 7.5; 150mM NaCl; 0.1% sodium deoxycholate; 1% Triton X-100; 5mM CaCl2; Protease Inhibitor Cocktail; 5mM sodium butyrate) for 10min on ice. Lysis Buffer with MNase (62μl) was then added for chromatin digestion and incubated at 37°C for exactly 10min. Then, 20mM EGTA was added to stop the reaction, followed by 13000rpm centrifugation for 5min at 4°C to sediment undigested debris. Supernatant was then transferred and equal amount of STOP buffer (50mM Tris-HCl, pH 7.5; 150mM NaCl; 0.1% sodium deoxycholate; 1% Triton X-100; 30mM EGTA; 30mM EDTA; Protease Inhibitor Cocktail; 5mM sodium butyrate) was added to the samples, which were always kept on ice.

Lysate (5μl) was mixed with 45μl of ProteinaseK (ProtK) Digestion Buffer (20mM HEPES; 1mM EDTA; 0.5% SDS) and incubated at 56°C for 30min. AMPure XP beads (50μl) were mixed with the digested lysate with 60μl of 20% PEG8000 1.25M NaCl for 15min at RT. Beads were separated on a magnet and washed twice with 80% Ethanol for 30 seconds. DNA was eluted in 12μl of Low-EDTA TE and measured using Qubit DNA High-Sensitivity kit to normalize lysate concentration between samples. DNA isolated in this step was used for the input sample. The volume of each undigested lysate was adjusted for equal concentration to obtain 1ml per IP using a 1:1 mix of Lysis Buffer and STOP Buffer.

Protein-A Dynabeads (10μl/IP) were washed twice in Blocking Buffer (0.5% BSA; 0.5% Tween in PBS) before being resuspended in Blocking buffer and coated with H3K27me3 [1μg/IP] (Cell Signalling, Cat#9733S) or H2AK119ub [0.4μg/IP] (Cell Signalling, Cat# 8240S) antibodies for 4 hours at 4°C. Once coated beads were magnet-separated and resuspended in 1ml of concentration-adjusted lysate. Samples were left rotating overnight at 4°C.

In the following day beads were magnet-separated and washed quickly with ice-cold washing buffers with Low Salt Buffer (0.1% SDS; 1% TritonX-100; 2mM EDTA; 20mM Tris-HCl, pH 8.1; 150mM NaCl; 0.1% sodium deoxycholate). IPs were then washed four times with Low Salt Buffer, twice with High Salt Buffer (0.1% SDS; 1% TritonX-100; 2mM EDTA; 20mM Tris-HCl, pH 8.1; 360mM NaCl; 0.1% sodium deoxycholate) and twice with LiCl buffer (0.25M LiCl; 1% NP40; 1.1% sodium deoxycholate; 1mM EDTA; 10mM Tris-HCl pH 8.1). Prior to elution all samples were rinsed once in TE. ChIP-DNA was eluted in ProtK-Digestion buffer by incubating at 56°C for 15min. Beads were separated and the supernatant was further digested for more 2 hours at 56°C. DNA was isolated using AMPure XP beads as described for the input sample.

For each nChIP-seq, 0.5μl of each sample was used for qPCR validation of enrichment at control regions (data not shown). 0.5μl of input samples were also used to verify the digestion efficiency using D1000 tapestation. Remaining DNA concentration was adjusted and used for library preparation using Ovation^®^ Ultralow Library System V2 following suppliers protocol. Amplified libraries were size-selected for dinucleotide fraction (350-600 bp fragments) using agarose gel-separation and MinElute Gel Extraction Kit (Qiagen). Sample quality was inspected using D1000 tapestation. Samples were sequenced with HiSeq2500 using single-end 50bp mode.

Adapters and low quality bases (<Q20) have been removed from the sequencing data with TrimGalore (v0.4.0) (Krueger) and Cutadapt (1.8.2) (Martin et al., 2011). Reads were then mapped to the mm10 genome with Bowtie2 (2.2.5) with options [--end-to-end-N1-q] (Langmead and Salzberg, 2012). Duplicates were discarded with Picard MarkDuplicates (1.65) with options [REMOVE_DUPLICATES=true] (DePristo et al., 2011), reads mapped with low quality (< q10) were removed using samtools (1.3) (Banito et al., 2009) and reads mapped on blacklisted regions from Encode Consortium (Consortium, 2012) were discarded. Bigwig files were created with bedtools genomeCoverageBed (2.25.0) (Quinlan and Hall, 2010), using a scale factor calculated on the total library (10.000.000/total reads) and loaded on UCSC genome browser (Kent et al., 2002).

ChIP-seq signal was then analyzed per window. A global analysis was first done on fixed windows (10Kb) spanning the whole genome, then on different genomic subcategories: active gene bodies, active promoters and intergenic regions. Active genes were defined in our previous study (Zylicz, Bousard et al., *in press*) as genes with a transcript having its TSS (refFlat annotation (Tyner et al., 2017) overlapping a peak of H3K27ac in noDOX samples. For genes having several active transcripts detected, the active gene was defined as starting at the minimum start of transcripts, and ending at the maximum end of transcripts. This way, 6096 active genes were defined genome-wide, 286 being on the X chromosome. The active gene bodies were defined as those active genes excluding the 2 first Kb downstream from TSS. Active promoters were defined as +/− 2Kb windows around the TSS of active genes. Intergenic regions were defined as 10Kb windows not overlapping a gene (active or inactive) and its promoter (2Kb downstream) or a peak of H3K27ac (Zylicz, Bousard et al., *in press*). Reads overlapping defined windows were then counted with featureCounts (1.5.1) (Liao et al., 2014) with default options.

For global analysis, counts normalization was performed based on counts falling in autosomal consensus peaks. For each histone mark, peaks were first identified in each sample using MACS2 (Zhang et al., 2008), with options [--broad-B-broad-cutoff 0.01] and with input as control, and only peaks with a minimum fold change of 3 were selected. Then, consensus peaks were defined as common regions between peaks identified in a minimum of 2 among the 4 noDOX samples using bedtools multiIntersectBed (2.25.0) and bedtools merge (2.25.0) (Quinlan and Hall, 2010). For each sample, a normalization factor was calculated with the trimmed mean of M-values method (TMM) from edgeR package (Fink et al., 2017), based on reads overlapping consensus peaks located on autosomes. To correct for chromatin accessibility or mappability bias, 10Kb windows with outliers counts in the input (counts superior or inferior to mean +/-1.5 sd – standard deviation) were discarded from the analysis. Moreover, to represent read accumulation between DOX and noDOX conditions, normalized initial counts from noDOX samples were subtracted to corresponding DOX normalized counts.

For genomic subcategories analysis (active gene bodies, active promoters and intergenic regions), windows that had less than one read per 50bp for more than 2 among the 8 samples were removed for the analysis. Normalization factors were calculated based on windows located on autosomes, with TMM method using edgeR (Fink et al., 2017). Linear regression was then fitted for each window according the following model: Y = clones + clones:condition, with Y being the log2(cpm) and condition being noDOX or DOX, using Voom function from Limma R package (Ritchie et al., 2015). Significance of coefficients was assessed by a moderated t-statistics and p-values were corrected by Benjamini-Hochberg procedure. Because of the high variability in proportion of cells with *Xist* induction, we quantified the number of cells with a *Xist* cloud by RNA FISH experiments: *Xist* FL DOX#1 - 46.64%, *Xist* FL-DOX#2 59.44%, *Xist* ΔB+C DOX#1 - 66.30%, *Xist* ΔB+C DOX#2 - 56.29%. Linear regression including the percentage of induction calculated by RNA FISH was also fitted for each window according the following model: ~0 + clones + clones:induction, using Voom function from Limma R package (Ritchie et al., 2015). The slope of this regression represents then the logFC between noDOX and DOX conditions if the induction of the cell population was complete (corrected logFC).

Metaplots were created using DeepTools (3.0.2) (Ramirez et al., 2014). Bigwigs of log2(FC) between DOX and noDOX samples were first created with personalized scaling according to normalized factors calculated above for active promoters using DeepTools bamCompare. Then, bigwigs of mean of log2(FC) between replicates were then created using DeepTools bigwigCompare with options [--binSize 100 --operation mean], matrix counts were then generated using DeepTools computeMatrix around TSS of active genes coordinates (see above) on X chromosome and autosomes separately and plots were then created using DeepTools PlotProfile.

### RNA-seq

Duplicates samples of *Xist* FL, *Xist* ΔA for *Xist* ΔB+C ES cells were differentiated until day 2 in DOX and noDOX conditions. Total RNA was isolated using NYZol (NZYTech) and then DNAse I treated (Roche) to remove contaminating DNA following the manufacturer’s recommendations. Initial RNA quality was checked by electrophoresis and sent to NOVOGENE for sequencing. Quality of the samples were verified on a 2100 Agilent Bioanalyser system and only samples with RIN score above 9 were processed. RNA (1μg) was used for 250-300bp insert cDNA library following manufacturer’s recommendations (except for *Xist* ΔB+C DOX#2, which only 100 ng of RNA was used for the library preparation using a low input method). Libraries were sequenced with NovaSeq 6000 platform using paired-end 150bp mode.

Reads were mapped on mm10 genome with Tophat (2.1.0) (Trapnell et al., 2009), with options [-g 1-x 1-N 5 --read-edit-dist 5 --no-coverage-search], with refFlat annotation (Tyner et al., 2017). Reads covering exons of each gene were then counted with featureCounts (1.5.1) with options [-C -p] (Liao et al., 2014). Bigwig files were created with Deeptools bamCoverage (2.2.4) (Ramirez et al., 2014), with option [--normalizeUsingRPKM] and loaded on UCSC genome browser (Kent et al., 2002).

Clustering of samples based on normalized counts of X-chromosome (calculated with cpm function from edgeR) was done with hclust function with parameter [method=‘Ward.D’], using Pearson correlation as distance.

Differential analysis was done on genes for which 6 among the 12 samples have a TPM superior to 1. Counts normalization was done based on counts falling in expressed autosomal genes, with the trimmed mean of M-values method (TMM) from edgeR package (Fink et al., 2017). Such as for nChIP-seq analysis, linear regression was then fitted for each gene with models including DOX/noDOX information, or percentage of induction calculated by RNA FISH (*Xist* FL DOX#1 - 46.6%, *Xist* FL DOX#2 - 51.7%, *Xist* ΔA DOX#1 - 53.2%, *Xist* ΔA DOX #2 - 54.5%, *Xist* ΔB+C DOX#1 - 61.4%, *Xist* ΔB+C DOX#2 - 56.7%).

Expression and ChIP-seq data were integrated as follow. First, for each mutant analyzed in both sets of data (*Xist* FL and *Xist* ΔB+C), four groups of genes were defined based on log2(FC) of RNA-seq: [−∞, −1.5], [−1.5, −1], [−1, −0.5] and [−0.5, ∞]. For each of these groups, the accumulation of normalized reads from ChIP-seq data at promoters of corresponding genes (DOX-noDOX signal, for genes with enough coverage (see ChIP-seq part above) was extracted, for each histone mark separately (H3K27me3, H2AK119ub). For each group of genes, the normalized ChIP-seq reads enrichment relative to noDOX between both cell lines was then compared using with a Wilcoxon test (Figure 4 - figure supplement 1F).

For each mark, promoters were divided in 2 categories: the ones with no accumulation or residual accumulation of H3K27me3 and H2AK119ub marks, and the ones with substantial accumulation of these repressive marks. The threshold between those two categories was defined as the mean + standard deviation (sd) of normalized signal accumulation between DOX and noDOX conditions (normalized reads DOX - normalized reads noDOX) on autosomes in *Xist* ΔB+C samples. Based on the data, the thresholds were 224 for H3K27me3 and 100 for H2AK119ub. Then, two categories of active promoters were defined based on both repressive marks: the ones with no or little accumulation for any of the 2 repressive marks (H3K27me3; H2AK119ub) and the ones with substantial accumulation of one or both repressive marks (Figure 4 - figure supplement 1G). The CpG content of each category was calculating using bedtools (2.25.0) with options “nuc --pattern G” and options “nuc --pattern CG”, and both were compared using a Wilcoxon test (Figure 4 – figure supplement 1H). Then, for each cell line (*Xist* FL, *Xist* ΔB+C), the expression log2(FC) of the genes not accumulating (or accumulating residual marks) and genes accumulating repressive marks in *Xist* ΔB+C were compared using Wilcoxon test. Moreover, inside a same category, expression log2(FC) was compared between *Xist* FL and *Xist* ΔB+C cell lines using a paired Wilcoxon test (Figure 4 F).

## Supporting information

## Acknowledgements

We thank Elphège Nora, Rafael Galupa, Martin Escamilla-Del-Arenal, Sérgio de Almeida and Cláudia Gil for critical reading of the manuscript. We are also grateful to members of the Heard lab (Curie Institute), members of the Carmo-Fonseca’s lab (iMM JLA), in particular to Kenny Rebelo, and also to Pierre Gestraud and Nicolas Servant (U900, Curie Institute) and Aurélie Teissandier (U934/UMR3215, Curie Institute) for their help and critical input to this project. We also want to thank the Bioimaging facility at iMM JLA for their technical support with fluorescent light microscopy.

This work was supported by Fundação para a Ciência e Tecnologia (FCT), project grants PTDC/BEX-BCM/2612/2014 (A.C.R. and S.T.d.R) and IF/00242/2014 (V.P. and S.T.d.R), by an ERC Advanced Investigator award ERC-ADG-677 2014 671027 attributed to E.H. and Sir Henry Wellcome Postdoctoral Fellowship (J.J.Z.), by the Scleroderma Research Foundation (Y.Q., H.Y.C) and by US National Institutes of Health NIH P50-HG007735 (H.Y.C.). H.Y.C. is an Investigator of the Howard Hughes Medical Institute.

## References

Almeida, M., Pintacuda, G., Masui, O., Koseki, Y., Gdula, M., Cerase, A., Brown, D., Mould, A., Innocent, C., Nakayama, M., et al. (2017). PCGF3/5-PRC1 initiates Polycomb recruitment in X chromosome inactivation. Science 356, 1081–1084.

Banito, A., Rashid, S.T., Acosta, J.C., Li, S., Pereira, C.F., Geti, I., Pinho, S., Silva, J.C., Azuara, V., Walsh, M., et al. (2009). Senescence impairs successful reprogramming to pluripotent stem cells. Genes Dev 23, 2134–2139.

Brockdorff, N. (2013). Noncoding RNA and Polycomb recruitment. RNA 19, 429–442.

Brown, S.D. (1991). XIST and the mapping of the X chromosome inactivation centre. Bioessays 13, 607–612.

Chaumeil, J., Augui, S., Chow, J.C., and Heard, E. (2008). Combined immunofluorescence, RNA fluorescent in situ hybridization, and DNA fluorescent in situ hybridization to study chromatin changes, transcriptional activity, nuclear organization, and X-chromosome inactivation. Methods Mol Biol 463, 297–308.

Chu, C., Zhang, Q.C., da Rocha, S.T., Flynn, R.A., Bharadwaj, M., Calabrese, J.M., Magnuson, T., Heard, E., and Chang, H.Y. (2015). Systematic discovery of Xist RNA binding proteins. Cell 161, 404–416.

Cirillo, D., Blanco, M., Armaos, A., Buness, A., Avner, P., Guttman, M., Cerase, A., and Tartaglia, G.G. (2016). Quantitative predictions of protein interactions with long noncoding RNAs. Nat Methods 14, 5–6.

Consortium, E.P. (2012). An integrated encyclopedia of DNA elements in the human genome. Nature 489, 57–74.

Cooper, S., Grijzenhout, A., Underwood, E., Ancelin, K., Zhang, T., Nesterova, T.B., Anil-Kirmizitas, B., Bassett, A., Kooistra, S.M., Agger, K., et al. (2016). Jarid2 binds mono-ubiquitylated H2A lysine 119 to mediate crosstalk between Polycomb complexes PRC1 and PRC2. Nat Commun 7, 13661.

da Rocha, S.T., Boeva, V., Escamilla-Del-Arenal, M., Ancelin, K., Granier, C., Matias, N.R., Sanulli, S., Chow, J., Schulz, E., Picard, C., et al. (2014). Jarid2 Is Implicated in the Initial Xist-Induced Targeting of PRC2 to the Inactive X Chromosome. Molecular cell 53, 301–316.

da Rocha, S.T., and Heard, E. (2017). Novel players in X inactivation: insights into Xist-mediated gene silencing and chromosome conformation. Nat Struct Mol Biol 24, 197–204.

Davidovich, C., Wang, X., Cifuentes-Rojas, C., Goodrich, K.J., Gooding, A.R., Lee, J.T., and Cech, T.R. (2015). Toward a consensus on the binding specificity and promiscuity of PRC2 for RNA. Molecular cell 57, 552–558.

Davidovich, C., Zheng, L., Goodrich, K.J., and Cech, T.R. (2013). Promiscuous RNA binding by Polycomb repressive complex 2. Nat Struct Mol Biol 20, 1250–1257.

de Napoles, M., Mermoud, J.E., Wakao, R., Tang, Y.A., Endoh, M., Appanah, R., Nesterova, T.B., Silva, J., Otte, A.P., Vidal, M., et al. (2004). Polycomb group proteins Ring1A/B link ubiquitylation of histone H2A to heritable gene silencing and X inactivation. Dev Cell 7, 663–676.

DePristo, M.A., Banks, E., Poplin, R., Garimella, K.V., Maguire, J.R., Hartl, C., Philippakis, A.A., del Angel, G., Rivas, M.A., Hanna, M., et al. (2011). A framework for variation discovery and genotyping using next-generation DNA sequencing data. Nature genetics 43, 491–498.

Engreitz, J.M., Pandya-Jones, A., McDonel, P., Shishkin, A., Sirokman, K., Surka, C., Kadri, S., Xing, J., Goren, A., Lander, E.S., et al. (2013). The Xist lncRNA exploits three-dimensional genome architecture to spread across the X chromosome. Science 341, 1237973.

Escamilla-Del-Arenal, M., da Rocha, S.T., and Heard, E. (2011). Evolutionary diversity and developmental regulation of X-chromosome inactivation. Hum Genet 130, 307–327.

Fink, J.J., Robinson, T.M., Germain, N.D., Sirois, C.L., Bolduc, K.A., Ward, A.J., Rigo, F., Chamberlain, S.J., and Levine, E.S. (2017). Disrupted neuronal maturation in Angelman syndrome-derived induced pluripotent stem cells. Nat Commun 8, 15038.

Ha, N., Lai, L.T., Chelliah, R., Zhen, Y., Yi Vanessa, S.P., Lai, S.K., Li, H.Y., Ludwig, A., Sandin, S., Chen, L., et al. (2018). Live-Cell Imaging and Functional Dissection of Xist RNA Reveal Mechanisms of X Chromosome Inactivation and Reactivation. iScience 8, 1–14.

Hoki, Y., Kimura, N., Kanbayashi, M., Amakawa, Y., Ohhata, T., Sasaki, H., and Sado, T. (2009). A proximal conserved repeat in the Xist gene is essential as a genomic element for X-inactivation in mouse. Development 136, 139–146.

Kalantry, S., and Magnuson, T. (2006). The Polycomb group protein EED is dispensable for the initiation of random X-chromosome inactivation. PLoS Genet 2, e66.

Kalantry, S., Mills, K.C., Yee, D., Otte, A.P., Panning, B., and Magnuson, T. (2006). The Polycomb group protein Eed protects the inactive X-chromosome from differentiation-induced reactivation. Nat Cell Biol 8, 195–202.

Kent, W.J., Sugnet, C.W., Furey, T.S., Roskin, K.M., Pringle, T.H., Zahler, A.M., and Haussler, D. (2002). The human genome browser at UCSC. Genome Res 12, 996–1006.

Krueger, F. Trim Galore! (http://www.bioinformatics.babraham.ac.uk/projects/trim_galore/).

Langmead, B., and Salzberg, S.L. (2012). Fast gapped-read alignment with Bowtie 2. Nat Methods 9, 357–359.

Leeb, M., and Wutz, A. (2007). Ring1B is crucial for the regulation of developmental control genes and PRC1 proteins but not X inactivation in embryonic cells. J Cell Biol 178, 219–229.

Liao, Y., Smyth, G.K., and Shi, W. (2014). featureCounts: an efficient general purpose program for assigning sequence reads to genomic features. Bioinformatics 30, 923–930.

Loda, A., Brandsma, J.H., Vassilev, I., Servant, N., Loos, F., Amirnasr, A., Splinter, E., Barillot, E., Poot, R.A., Heard, E., et al. (2017). Genetic and epigenetic features direct differential efficiency of Xist-mediated silencing at X-chromosomal and autosomal locations. Nat Commun 8, 690.

Lu, Z., Zhang, Q.C., Lee, B., Flynn, R.A., Smith, M.A., Robinson, J.T., Davidovich, C., Gooding, A.R., Goodrich, K.J., Mattick, J.S., et al. (2016). RNA Duplex Map in Living Cells Reveals Higher-Order Transcriptome Structure. Cell 165, 1267–1279.

Lv, Q., Yuan, L., Song, Y., Sui, T., Li, Z., and Lai, L. (2016). D-repeat in the XIST gene is required for X chromosome inactivation. RNA Biol 13, 172–176.

Maenner, S., Blaud, M., Fouillen, L., Savoye, A., Marchand, V., Dubois, A., Sanglier-Cianferani, S., Van Dorsselaer, A., Clerc, P., Avner, P., et al. (2010). 2-D structure of the A region of Xist RNA and its implication for PRC2 association. PLoS Biol 8, e1000276.

Martin, M. (2011). Cutadapt removes adapter sequences from high-throughput sequencing 822 reads. 2011 17.

McHugh, C.A., Chen, C.K., Chow, A., Surka, C.F., Tran, C., McDonel, P., Pandya-Jones, A., Blanco, M., Burghard, C., Moradian, A., et al. (2015). The Xist lncRNA interacts directly with SHARP to silence transcription through HDAC3. Nature 521, 232–236.

Mendenhall, E.M., Koche, R.P., Truong, T., Zhou, V.W., Issac, B., Chi, A.S., Ku, M., and Bernstein, B.E. (2010). GC-rich sequence elements recruit PRC2 in mammalian ES cells. PLoS Genet 6, e1001244.

Minajigi, A., Froberg, J.E., Wei, C., Sunwoo, H., Kesner, B., Colognori, D., Lessing, D., Payer, B., Boukhali, M., Haas, W., et al. (2015). Chromosomes. A comprehensive Xist interactome reveals cohesin repulsion and an RNA-directed chromosome conformation. Science 349.

Moindrot, B., Cerase, A., Coker, H., Masui, O., Grijzenhout, A., Pintacuda, G., Schermelleh, L., Nesterova, T.B., and Brockdorff, N. (2015). A Pooled shRNA Screen Identifies Rbm15, Spen, and Wtap as Factors Required for Xist RNA-Mediated Silencing. Cell Rep 12, 562–572.

Monfort, A., Di Minin, G., Postlmayr, A., Freimann, R., Arieti, F., Thore, S., and Wutz, A. (2015). Identification of Spen as a Crucial Factor for Xist Function through Forward Genetic Screening in Haploid Embryonic Stem Cells. Cell Rep 12, 554–561.

Nesterova, T.B., Slobodyanyuk, S.Y., Elisaphenko, E.A., Shevchenko, A.I., Johnston, C., Pavlova, M.E., Rogozin, I.B., Kolesnikov, N.N., Brockdorff, N., and Zakian, S.M. (2001). Characterization of the genomic Xist locus in rodents reveals conservation of overall gene structure and tandem repeats but rapid evolution of unique sequence. Genome Res 11, 833–849.

Patil, D.P., Chen, C.K., Pickering, B.F., Chow, A., Jackson, C., Guttman, M., and Jaffrey, S.R. (2016). m(6)A RNA methylation promotes XIST-mediated transcriptional repression. Nature 537, 369–373.

Pintacuda, G., Wei, G., Roustan, C., Kirmizitas, B.A., Solcan, N., Cerase, A., Castello, A., Mohammed, S., Moindrot, B., Nesterova, T.B., et al. (2017). hnRNPK Recruits PCGF3/5-PRC1 to the Xist RNA B-Repeat to Establish Polycomb-Mediated Chromosomal Silencing. Molecular cell 68, 955–969 e910.

Plath, K., Fang, J., Mlynarczyk-Evans, S.K., Cao, R., Worringer, K.A., Wang, H., de la Cruz, C.C., Otte, A.P., Panning, B., and Zhang, Y. (2003). Role of histone H3 lysine 27 methylation in X inactivation. Science 300, 131–135.

Quinlan, A.R., and Hall, I.M. (2010). BEDTools: a flexible suite of utilities for comparing genomic features. Bioinformatics 26, 841–842.

Ramirez, F., Dundar, F., Diehl, S., Gruning, B.A., and Manke, T. (2014). deepTools: a flexible platform for exploring deep-sequencing data. Nucleic Acids Res 42, W187–191.

Ridings-Figueroa, R., Stewart, E.R., Nesterova, T.B., Coker, H., Pintacuda, G., Godwin, J., Wilson, R., Haslam, A., Lilley, F., Ruigrok, R., et al. (2017). The nuclear matrix protein CIZ1 facilitates localization of Xist RNA to the inactive X-chromosome territory. Genes Dev 31, 876–888.

Riising, E.M., Comet, I., Leblanc, B., Wu, X., Johansen, J.V., and Helin, K. (2014). Gene silencing triggers polycomb repressive complex 2 recruitment to CpG islands genome wide. Molecular cell 55, 347–360.

Ritchie, M.E., Phipson, B., Wu, D., Hu, Y., Law, C.W., Shi, W., and Smyth, G.K. (2015). limma powers differential expression analyses for RNA-sequencing and microarray studies. Nucleic Acids Res 43, e47.

Rutenberg-Schoenberg, M., Sexton, A.N., and Simon, M.D. (2016). The Properties of Long Noncoding RNAs That Regulate Chromatin. Annu Rev Genomics Hum Genet 17, 69–94.

Sarma, K., Levasseur, P., Aristarkhov, A., and Lee, J.T. (2010). Locked nucleic acids (LNAs) reveal sequence requirements and kinetics of Xist RNA localization to the X chromosome. Proc Natl Acad Sci U S A 107, 22196–22201.

Schindelin, J., Arganda-Carreras, I., Frise, E., Kaynig, V., Longair, M., Pietzsch, T., Preibisch, S., Rueden, C., Saalfeld, S., Schmid, B., et al. (2012). Fiji: an open-source platform for biological-image analysis. Nat Methods 9, 676–682.

Schmitz, S.U., Grote, P., and Herrmann, B.G. (2016). Mechanisms of long noncoding RNA function in development and disease. Cell Mol Life Sci 73, 2491–2509.

Senner, C.E., Nesterova, T.B., Norton, S., Dewchand, H., Godwin, J., Mak, W., and Brockdorff, N. (2011). Disruption of a conserved region of Xist exon 1 impairs Xist RNA localisation and X-linked gene silencing during random and imprinted X chromosome inactivation. Development 138, 1541–1550.

Silva, J., Mak, W., Zvetkova, I., Appanah, R., Nesterova, T.B., Webster, Z., Peters, A.H., Jenuwein, T., Otte, A.P., and Brockdorff, N. (2003). Establishment of histone h3 methylation on the inactive X chromosome requires transient recruitment of Eed-Enx1 polycomb group complexes. Dev Cell 4, 481–495.

Sunwoo, H., Colognori, D., Froberg, J.E., Jeon, Y., and Lee, J.T. (2017). Repeat E anchors Xist RNA to the inactive X chromosomal compartment through CDKN1A-interacting protein (CIZ1). Proc Natl Acad Sci U S A 114, 10654–10659.

Tang, Y.A., Huntley, D., Montana, G., Cerase, A., Nesterova, T.B., and Brockdorff, N. (2010). Efficiency of Xist-mediated silencing on autosomes is linked to chromosomal domain organisation. Epigenetics Chromatin 3, 10.

Tavares, L., Dimitrova, E., Oxley, D., Webster, J., Poot, R., Demmers, J., Bezstarosti, K., Taylor, S., Ura, H., Koide, H., et al. (2012). RYBP-PRC1 complexes mediate H2A ubiquitylation at polycomb target sites independently of PRC2 and H3K27me3. Cell 148, 664–678.

Trapnell, C., Pachter, L., and Salzberg, S.L. (2009). TopHat: discovering splice junctions with RNA-Seq. Bioinformatics 25, 1105–1111.

Tyner, C., Barber, G.P., Casper, J., Clawson, H., Diekhans, M., Eisenhart, C., Fischer, C.M., Gibson, D., Gonzalez, J.N., Guruvadoo, L., et al. (2017). The UCSC Genome Browser database: 2017 update. Nucleic Acids Res 45, D626–D634.

Wang, J., Mager, J., Chen, Y., Schneider, E., Cross, J.C., Nagy, A., and Magnuson, T. (2001). Imprinted X inactivation maintained by a mouse Polycomb group gene. Nature genetics 28, 371–375.

Wutz, A., Rasmussen, T.P., and Jaenisch, R. (2002). Chromosomal silencing and localization are mediated by different domains of Xist RNA. Nature genetics 30, 167–174.

Yamada, N., Hasegawa, Y., Yue, M., Hamada, T., Nakagawa, S., and Ogawa, Y. (2015). Xist Exon 7 Contributes to the Stable Localization of Xist RNA on the Inactive X-Chromosome. PLoS Genet 11, e1005430.

Yen, Z.C., Meyer, I.M., Karalic, S., and Brown, C.J. (2007). A cross-species comparison of X-chromosome inactivation in Eutheria. Genomics 90, 453–463.

Zhang, Y., Liu, T., Meyer, C.A., Eeckhoute, J., Johnson, D.S., Bernstein, B.E., Nusbaum, C., Myers, R.M., Brown, M., Li, W., et al. (2008). Model-based analysis of ChIP-Seq (MACS). Genome Biol 9, R137.

Zhao, J., Sun, B.K., Erwin, J.A., Song, J.J., and Lee, J.T. (2008). Polycomb proteins targeted by a short repeat RNA to the mouse X chromosome. Science 322, 750–756.

