## Supplementary material for "Exploring the role of Polycomb recruitment in *Xist*-mediated silencing of the X chromosome in ES cells"

Includes:

- Figure 1 - source data 1
- Figure 1 - source data 2
- Figure 2 –source data 1 (only the legend; this file is the excel format)
- Figure 4 – source data 1
- Supplementary File 1
- Supplementary File 2
- Figure 1 –figure supplement 1
- Figure 1 –figure supplement 2
- Figure 2 –figure supplement 1
- Figure 3 –figure supplement 1
- Figure 3 –figure supplement 2
- Figure 4 – figure supplement 1

| *Xist* mutant | Clone Number | Size of deletion | Deleted region | % of cells with *Xist* domains | *Xist* coating | Nº of experiments | Total cell counted |
|---|---|---|---|---|---|---|---|
| *Xist* FL | n.a. | n.a. | n.a. | 45 ± 6 % | +++++ | 3 | 956 |
| *Xist* ΔA | n.a. | 917 bp | Δ1-917 | 53 ± 9 % | ++++ | 3 | 990 |
| *Xist* ΔF+B+C | 1 | 3,908 bp | Δ890-4796 | 77 ± 4 % | +++ | 3 | 811 |
| | 2 | 3,908 bp | Δ890-4796 | n.d. | +++ | n.d. | n.a. |
| *Xist* ΔF+B | 1 | 2,119 bp | Δ875-2982 | 76 ± 3 % | +++ | 3 | 869 |
| | 2 | 2,130 bp | Δ862-2990 | 87% | +++ | 1 | 272 |
| *Xist* ΔB+C | 1 | 2,109 bp | Δ2,703-4810 | 60 ± 8 % | +++++ | 3 | 952 |
| | 2 | 2,049 bp | Δ2,699-4744 ( + 45 bp insertion) | 7% | +++++ | 1 | 269 |
| *Xist* ΔB+1/2C* | 1 | 1,383 bp | Δ2,703-4,084 | 60 ± 3 % | +++++ | 3 | 921 |
| | 2 | 454 bp & 810 bp | Δ2,705-3,158 & Δ3,558-4,367 | 52% | +++++ | 1 | 298 |
| *Xist* ΔB | 1 | 335 bp | Δ2,707-3,041 | 67 ± 4 % | +++++ | 3 | 978 |
| | 2 | 337 bp | Δ2,705-3,041 | 44% | +++++ | 1 | 365 |
| *Xist* ΔC | 1 | 1,716 bp | Δ3,041-4,752 | 71 ± 2 % | +++++ | 3 | 863 |
| | 2 | 1,716 bp | Δ3,041-4,752 | 34% | +++++ | 1 | 298 |

**Figure 1 - source data 1 – Full set of *Xist*-TetOP mutants generated**

Summary table of the analysis of full set of the *Xist*-TetOP mutants generated (including the second clone per type of mutation) in terms of: deletion size, coordinates of the deleted regions (based on Ensembl *Xist* exonic sequence); number of cells with *Xist* domains (in % ± S.E.M.) and capacity of *Xist* coating (+++++ indicates *Xist* cloud signals similar to *Xist* FL; ++++ and +++ indicate *Xist* cloud signals of decreased size compared to *Xist* FL); n.d. – not done; n. a. – not applied; * To generate *Xist* ΔB+1/2C mutants, we use a 3'end gRNA which recognized a unique region within repeat C, however, several other sequences with only 1 or 2 mismatched can be found within this repeat and were likely to be targeted, creating distinct deletions as can be seen for the two clones of this type of mutant.

| *Xist* mutant | Clone | JARID2 | EZH2 | H3K27me3 | RING1B | H2AK119ub |
|---|---|---|---|---|---|---|
| *Xist* FL | n.a. | 88 ± 4 %<br>(4 exp, n = 275) | 85 ± 7 %<br>( 3 exp, n = 242) | 92 ± 7 %<br>(3 exp, n = 248) | 48 ± 14 %<br>(4 exp, n = 313) | 96 ± 1 %<br>(3 exp, n = 223) |
| *Xist* ΔA | n.a. | 78 ± 4 %<br>(3 exp, n = 261) | 78 ± 11 %<br>(3 exp, n = 243) | 97 ± 2 %<br>( 3 exp, n = 228) | 50 ± 10 %<br>(4 exp, n = 302) | 97 ± 2 %<br>(3 exp, n = 269) |
| *Xist* ΔF+B+C | 1 | 1 ± 1 %<br>(3 exp, n = 257) | 1 ± 1 %<br>(3 exp, n = 232) | 3 ± 2 %<br>(3 exp, n = 236) | 0 ± 0 %<br>(2 exp, n = 143) | 1 ± 0 %<br>(3 exp, n = 227) |
| | 2 | n.d. | n.d. | 0%<br>(1 exp, n = 85) | n.d. | 2%<br>(1 exp, n =83) |
| *Xist* ΔF+B | 1 | 1 ± 1 %<br>(2 exp, n = 204) | 2 ± 2 %<br>(2 exp, n = 180) | 18 ± 15 %<br>(2 exp, n = 198) | 0 ± 0 %<br>(3 exp, n = 218) | 28 ± 7 %<br>(2 exp, n = 154) |
| | 2 | 14%<br>(1 exp, n = 93) | 18%<br>(1 exp, n = 180) | 32%<br>(1 exp, n = 66) | 0%<br>(1 exp, n = 57) | 32%<br>(1 exp, n = 57) |
| *Xist* ΔB+C | 1 | 0 ± 0 %<br>(3 exp, n = 237) | 0 ± 0 %<br>(3 exp, n = 216) | 1 ± 0 %<br>(3 exp, n = 208) | 0 ± 0 %<br>(4 exp, n = 292) | 1 ± 1 %<br>(2 exp, n = 181) |
| | 2 | 0%<br>(1 exp, n = 68) | 0%<br>(1 exp, n = 63) | 5%<br>(1 exp, n = 63) | 0 ± 0 %<br>(2 exp, n = 98) | 3%<br>(1 exp, n = 65) |
| *Xist* ΔB+1/2C | 1 | 0 ± 0 %<br>(2 exp, n = 202) | 0 ± 0 %<br>(2 exp, n = 165) | 5 ± 0 %<br>(2 exp, n = 182) | 0%<br>(1 exp, n = 54) | 17 ± 2 %<br>(3 exp, n = 262) |
| | 2 | 1%<br>(1 exp, n = 87) | n.d. | 5%<br>(1 exp, n = 63) | 0%<br>(1 exp, n = 100) | 14%<br>(1 exp, n = 100) |
| *Xist* ΔB | 1 | 6 ± 3 %<br>(3 exp, n = 265) | 18 ± 5 %<br>(3 exp, n = 270) | 49 ± 7 %<br>(3 exp, n = 240) | 1 ± 1 %<br>(3 exp, n = 217) | 61 ± 8 %<br>(3 exp, n = 257) |
| | 2 | n.d. | n.d. | 38%<br>(1 exp, n = 90) | n.d. | 63 ± 16 %<br>(2 exp, n = 93) |
| *Xist* ΔC | 1 | 89 ± 4 %<br>(3 exp, n = 255) | 81 ± 9%<br>(2 exp, n = 172) | 78 ± 2 %<br>(3 exp, n = 195) | 37 ± 4 %<br>(3 exp, n = 240) | 99 ± 0 %<br>(3 exp, n = 288) |
| | 2 | 89%<br>(1 exp, n = 72) | n.d. | 65%<br>(1 exp, n = 63) | n.d. | 97%<br>(1 exp, n = 72) |

**Figure 1 - source data 2 - Summary table of IF/*Xist* RNA FISH experiments in the full set of *Xist*-TetOP mutants generated**

The table displays the percentage (± S.E.M.) of *Xist*-coated chromosomes exhibiting enrichment of the PcG proteins (JARID2, EZH2 and RING1B) and histone marks (H3K27me3 and H2AK119ub); a minimum of 50 *Xist*-coated chromosomes were counted per experiment (exp); n – number of cells counted in at least one or more biological replicates; n.a. – not applied; n.d. – not done.

**Figure 2 - source data 1 – Full list of ChIRP-MS peptide counts for *Xist* FL (noDOX and DOX) and *Xist* ΔB+C (DOX) – supplementary excel file**

<u>Sheet 1</u> – Selection of peptide counts for the 81 *Xist* hits [according to (Chu et al., 2015)] found in *Xist* FL and *Xist* ΔB+C interactome (with a minimum of 2.5 DOX/noDOX fold-change in *Xist* FL or *Xist* ΔB+C; Weakly annotated protein isoforms with an Annotation score in UniprotKB < 3 (out of 5) were excluded.
<u>Sheet 2</u> – Filtered list of total number of peptides counts for in *Xist* FL (noDOX and DOX) and *Xist* ΔB+C; In this list, only peptides with a minimum of 2.5 DOX/noDOX fold-change in *Xist* FL were selected; Weakly annotated protein isoforms with an Annotation score in UniprotKB < 3 (out of 5) were excluded; this was the case of two poorly annotated isoforms of hnRNPK: tr|Q3TL71, less bound to *Xist* ΔB+C and tr|Q3U6X2, equally bound to *Xist* FL and ΔB+C.
<u>Sheet 3</u> – Total number of peptide counts in *Xist* FL (noDOX and DOX) and *Xist* ΔB+C (DOX).

| *Xist* mutant | Clone Number | ***Lamp2*** **(D2)** | ***Pgk1*** **(D2)** | ***Pgk1*** **(D4)** | ***Rnf12*** **(D4)** |
|---|---|---|---|---|---|
| *Xist* FL (noDOX) | n.a. | 58 ± 9% (2 exp, n = 245) | 53 ± 7% (3 exp, n = 308) | 68 ± 8% (2 exp, n =353) | 73% (1 exp, n = 106) |
| *Xist* FL | n.a. | 10 ± 1 % (3 exp, n = 345) | 9 ± 3% (3 exp, n = 375) | 11 ± 2 % (4 exp, n = 363) | 19 ± 9 % (2 exp, n = 211) |
| *Xist* ΔA | n.a. | 75 ± 2 % (3 exp, n = 344) | 68 ± 6% (3 exp, n = 352) | 64 ± 7 % (3 exp, n = 332) | 85 ± 5 % (2 exp, n = 228) |
| *Xist* ΔF+B+C | 1 | 21 ±1 % (3 exp, n = 351) | 15 ± 1% (2 exp, n = 246) | 14 ± 2 % (3 exp, n = 353) | n.d. |
| | 2 | 10% (1 exp, n = 81) | n.d. | n.d. | n.d. |
| *Xist* ΔF+B | 1 | 5% (1 exp, n = 107) | 12 ± 2 % (2 exp, n = 216) | 21 ± 8 % (3 exp, n = 327) | 23 ± 4 % (2 exp, n = 210) |
| | 2 | n.d. | n.d. | 9% (1 exp, n =111) | 12% (1 exp, n = 105) |
| *Xist* ΔB+C | 1 | 17 ± 1 % (3 exp, n = 331) | 11 ± 1 % (3 exp, n = 367) | 17 ± 4 % (4 exp, n = 377) | 17% (1 exp, n = 97) |
| | 2 | 20% (1 exp, n = 66) | 16% (1 exp, n = 64) | n.d. | n.d. |
| *Xist* ΔB+1/2C | 1 | 18% (1 exp, n = 99) | 11 ± 2% (2 exp, n = 260) | 19 ± 1 % (2 exp, n = 246) | 5% (1 exp, n = 111) |
| | 2 | n.d. | n.d. | 6% (1 exp, n = 112) | n.d. |
| *Xist* ΔB | 1 | 13 ± 1% (3 exp, n = 330) | 11 ± 1% (2 exp, n = 220) | 14 ± 1 % ( 2 exp, n = 205) | n.d. |
| | 2 | n.d. | n.d. | 23% (1 exp, n = 93) | n.d. |
| *Xist* ΔC | 1 | 8± 2% (3 exp, n = 292) | 5 ± 2 % (2 exp, n = 198) | 5 ± 1 % ( 2 exp, n = 327) | n.d. |
| | 2 | n.d. | 0% (1 exp, n = 59) | n.d. | n.d. |

**Figure 4 - source data 1 - Summary table of combined *Xist* with X-linked nascent-transcript RNA FISH for *Pgk1, Lamp2* and *Rnf12* in the full set of *Xist*-TetOP mutants**

The table displays the percentage (± S.E.M.) of *Xist*-coated chromosomes exhibiting an active *Pgk1* (at D2 and D4), *Lamp2* (at D2) and *Rnf12* gene (at D4); a minimum of 50 *Xist*-coated chromosomes were counted per experiment (exp); for *Xist* FL noDOX a minimum of 100 cells (which do not have *Xist*-coated chromosome) were counted; n – number of cells counted in at least one or more biological replicate (exp); n.d. – not done; n.a. – not applied.

| *Xist* mutants | 5'end /3'end sgRNAs | Primers to confirm deletion (F/R) | Primers to confirm loss of WT sequence (F/R) |
|---|---|---|---|
| *Xist* ΔF+B+C | TCACGCAGAAGCCATAATGG/ CTTGAGAGATGATACCTCCA | CTGCTGATCGTTTGGTGCTG/ CAGACCTGTGTTTGCCCCTT | CTGCTGATCGTTTGGTGCTG/ ATCAAGGCGAATCCCGCAAC |
| *Xist* ΔF+B | TCACGCAGAAGCCATAATGG/ AGGGCTGGACTGGATTGGGT | TGGTGCTGTGTGAGTGAACC/ TTAGCACTGAATCAATGAAGA | CTGCTGATCGTTTGGTGCTG/ ATCAAGGCGAATCCCGCAAC |
| *Xist* ΔB+C | TATAACAGTAAGTCTGATAG/ GTGTATCTTGATTAACATGA | ATGACTGGATGTCAGGAGTA/ CAGACCTGTGTTTGCCCCTT | ATGACTGGATGTCAGGAGTA/ CTGAGTCTTGAGGAGAATCT |
| *Xist* ΔB+1/2C | TATAACAGTAAGTCTGATAG/ CATACTGACTTCTAGAGTCA* | ATGACTGGATGTCAGGAGTA/ CAGACCTGTGTTTGCCCCTT | ATGACTGGATGTCAGGAGTA/ CTGAGTCTTGAGGAGAATCT |
| *Xist* ΔB | TATAACAGTAAGTCTGATAG/ CTCTAAGTAGAAGTGGGCTT | ATGACTGGATGTCAGGAGTA/ TTAGCACTGAATCAATGAAGA | ATGACTGGATGTCAGGAGTA/ CTGAGTCTTGAGGAGAATCT |
| *Xist* ΔC | CTCTAAGTAGAAGTGGGCTT/ GTGTATCTTGATTAACATGA | CCAGGCCCAGATACTTTCAG/ CAGACCTGTGTTTGCCCCTT | TCCATGGACAAGTAAACAAAGAA/ TGTTTGCCCCTTTGCTAAAT |

**Supplementary File 1 – List of gRNA sequences and primers used for *CRISPR/Cas9* editing of the different *Xist*-TetOP mutants**

Primer sequences to confirm deletion and loss of wild-type (WT) allele are displayed for each *Xist* mutant; * highlights for the fact that the 3'end gRNA used to generate ΔB+1/2C was designed to a unique region within the repeat C, but several sequences with only 1 or 2 mismatched can be found within this repeat and are likely to be targeted to generate different types of ΔB+1/2C mutants.

| **RT-PCR analysis** | Sequences (F/R) |
|---|---|
| *Xist* exon 1-exon 3 | GCTGGTTCGTCTATCTTGTGGG / CAGAGTAGCGAGGACTTGAAGAG |
| *Xist* before repeat B | ATGACTGGATGTCAGGAGTA / CTGAGTCTTGAGGAGAATCT |
| *Xist* before repeat C | TCCATGGACAAGTAAACAAAGAA / TGTTTGCCCCTTTGCTAAAT |
| *Gapdh* | AACTTTGGCATTGTGGAAGG / ACACATTGGGGGTAGGAACA |

**Supplementary File 2- Primer sequences for RT-PCR analysis of *Xist* mutants (used in Figure 1 – supplement figure 1B)**

**Figure 1 - figure supplement 1 – Characterization of the novel *Xist*-TetOP mutants**

A. Deletion mapping by Sanger sequencing and expression analysis across deleted region in the novel *Xist* ΔF+B+C, ΔB+F, ΔB+C, ΔB+1/2C, ΔB and ΔC mutants (this analysis is for clone 1 of each mutant type); red arrows indicate forward primer and green arrows represent reverse primers; the primer of the left is the sequenced primer for each mutant; B means PCR blank.
B. RT-PCR analysis of the splicing pattern and expression across the repeat B and C regions of the different *Xist*-TetOP mutants; Primer pairs indicated in the scheme in green (see Materials and Methods); B means PCR blank.
C. *Xist* RNA FISH analysis upon D4 of differentiation in the presence of DOX (also noDOX for *Xist* FL) in the *Xist* ΔF+B+C, ΔB+F, ΔB+C, ΔB+1/2C, ΔB and ΔC mutants (this analysis is for clone 1 of each mutant type); values represent the % ± standard error (S.E.M.) of cells with a *Xist*-coated chromosome (at least 3 biological replicates with a minimum of 250 cells counted per replicate); Scale bar: 10 µm.

**Figure 1 - figure supplement 2 – Recruitment of a PRC2 member (EZH2), a PRC2 co-factor (JARID2) and PRC1 member (RING1B) in the different *Xist*-TetOP mutants**

A. Representative images of combined IF for JARID2 (green) with RNA FISH for *Xist* (red) in *Xist*-TetOP lines (for clone 1 of each mutant type) upon D2 in the presence of DOX; DAPI in blue.
B. Representative images of combined IF for RING1B (green) with RNA FISH for *Xist* (red) in *Xist*-TetOP lines (for clone 1 of each mutant type) upon D2 in the presence of DOX; DAPI in blue; Scale bar: 10 µm.
C. Graph representing the % of *Xist*-coated chromosomes enriched for JARID2, EZH2 and RING1B in the different *Xist*-TetOP mutants (for clone 1 of each mutant type) from to 2-to-4 independent experiments. A minimum of 50 *Xist*-coated chromosomes were counted per experiment. Significant differences from unpaired Student's *t*-test, comparing mutants to *Xist* FL are indicated as * p-value < 0.05.

**Figure 2 - figure supplement 1 – Quality check of ChIRP procedure in *Xist* FL and *Xist* ΔB+C cells**

A. RT-qPCR with three primer pairs along *Xist* to evaluate RNA retrieval after ChIRP procedure for *Xist* FL (in noDOX and DOX conditions) and *Xist* ΔB+C (DOX) at D3.
B. Table showing the percentage of cells exhibiting a *Xist*-coated X chromosome for *Xist* FL (both noDOX and DOX) and *Xist* ΔB+C (DOX) as determined by *Xist* RNA FISH used for ChIRP-MS; A minimum of 500 cells were counted.
C. Blot visualized with Coomassie blue staining showing the band pattern of proteins displayed by *Xist* FL (both noDOX and DOX) and *Xist* ΔB+C (DOX) after ChIRP.

**Figure 3 - figure supplement 1 – nChIP-seq confirms residual enrichment of H3K27m3 and H2AK119ub marks at X-linked active genes**

A. Normalized signal of H3K27me3 and H2AK119ub around *HoxC* cluster; shown is the signal of each sample around these cluster, normalized by the size of the library.
B. Barplot representing percentages of H3K27me3 and H2AK119ub reads mapping on X chromosome (chrX) in each sample.
C. Violin plots quantifying H3K27me3 and H2AK119ub enrichment over intergenic regions, active promoters and active gene bodies on X chromosome and on autosomes in *Xist* FL and *Xist* ΔB+C cell lines upon DOX induction at day of differentiation; Shown is the calculated log2 fold change of DOX vs noDOX conditions; n = indicates the number of genes analyzed; p-values were calculated using unilateral Wilcoxon test, comparing X chromosome (chrX) and autosomal enrichment of PcG marks for each genomic region.

**Figure 3 - figure supplement 2 – Normalization of H3K27me3 and H2AK119ub enrichment over the X chromosome in *Xist* FL and *Xist* ΔB+C based on *Xist* induction levels**

A. Table showing the percentage of cells exhibiting a *Xist*-coated X chromosome (chrX) for the different duplicates of *Xist* FL and *Xist* ΔB+C in DOX and noDOX conditions as determined by *Xist* RNA FISH; A minimum of 500 cells were counted to calculate the percentage of cells with a *Xist*-coated chrX.
B. Violin plots quantifying H3K27me3 and H2AK119ub enrichment over intergenic regions, active promoters and gene bodies on chrX in *Xist* FL and *Xist* ΔB+C upon DOX induction at day 2 of differentiation after normalization for the percentage of

cells with *Xist*-coated chromosomes. Shown is the log2 fold change of DOX vs noDOX conditions; n = indicates the number of genes analyzed; p-values were calculated using paired Wilcoxon test, comparing *Xist* FL and *Xist* ΔB+C cell lines.

**Figure 4 - figure supplement 1 –Assessment of transcriptional changes by RNA-seq in *Xist* FL, *Xist* ΔA and *Xist* ΔB+C induced cells**

A. Genome browsers plots showing RNA-seq reads on *Xist/Tsix* genes for *Xist* FL, *Xist* ΔA and *Xist* ΔB+C mutants in DOX and noDOX conditions at D2 of differentiation; yellow boxes display the deleted regions in both *Xist* ΔA and *Xist* ΔB+C.
B. Barplot representing percentages of RNA-seq reads mapping on X-chromosome (chrX) in each sample.
C. Table showing the percentage of cells exhibiting an *Xist*-coated chrX for the different duplicates of *Xist* FL, *Xist* ΔA and *Xist* ΔB+C in DOX and noDOX conditions as determined by *Xist* RNA FISH; at least 500 cells were counted to estimate the percentage of cells with a *Xist*-coated chrX.
D. Violin plots displaying the average log2(fold-change) in gene expression between DOX and noDOX conditions on chrX and autosomes in *Xist* FL, *Xist* ΔA and *Xist* ΔB+C at D2 after normalization for the percentage of cells with a *Xist*-coated chrX; n = indicates the number of genes analyzed; p-values for chrX were calculated using paired Wilcoxon test; n = indicates the number of genes analyzed.
E. Plots displays the comparison of log2(fold-change) in X-linked gene silencing upon DOX induction between *Xist* FL and *Xist* ΔB+C at D2 of differentiation; Limma *t*-test did not find any gene differentially expressed between *Xist* FL and *Xist* ΔB+C.
F. Boxplots displaying the normalized read enrichment at promoters for H3K27me3 and H2AK119ub upon DOX induction for distinct categories of X-linked genes with different degrees of gene silencing between DOX and noDOX conditions in both *Xist* FL and *Xist* ΔB+C at D2; p-values were calculated using Wilcoxon test; numbers inside the boxplots indicate the number of genes analyzed.
G. Boxplots displaying H3K27me3 and H2AK119ub normalized enrichment levels at promoters upon induction in two categories of X-linked genes: with no or little accumulation versus with accumulation of these PcG marks in induced *Xist* ΔB+C cells; p-values were calculated using Wilcoxon test; n = indicates the number of genes analyzed

H. CpG content in promoters of X-linked genes with no or little accumulation versus with accumulation of PcG marks in induced *Xist* ΔB+C cells; p-values were calculated using Wilcoxon test; Numbers inside the boxplots indicate the number of genes analyzed.

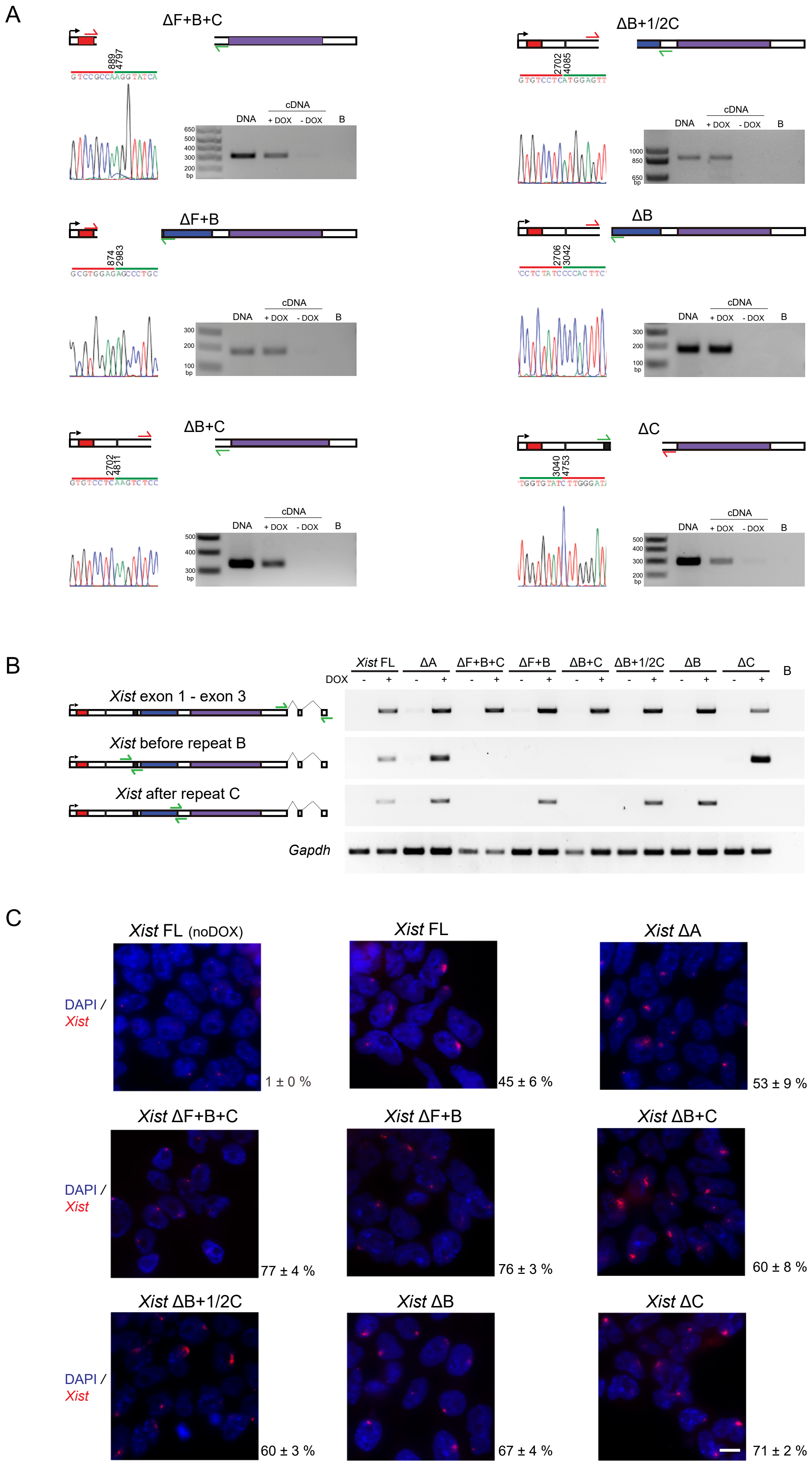


Figure 1 - figure supplement 1

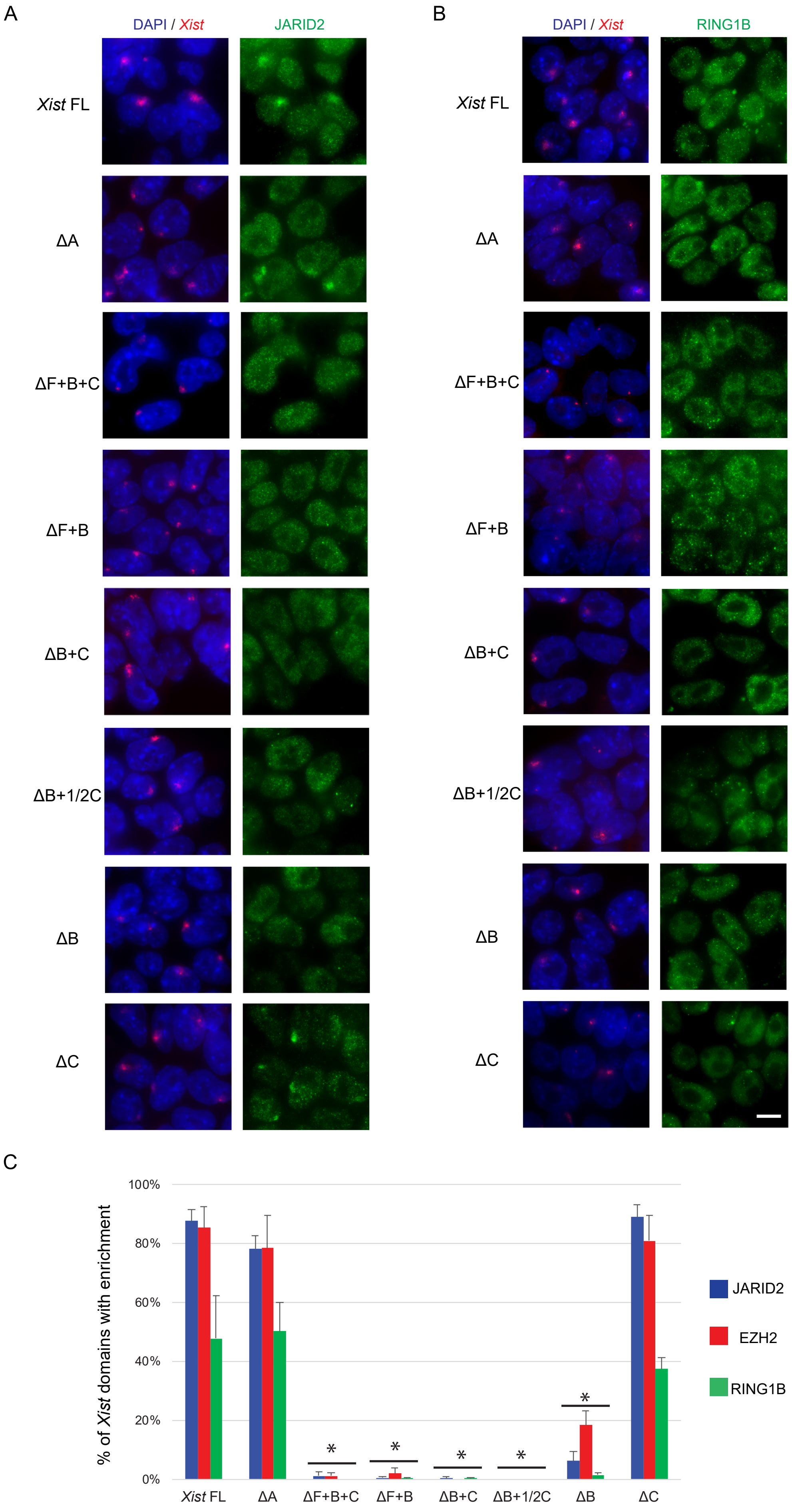


Figure 1 - figure supplement 2

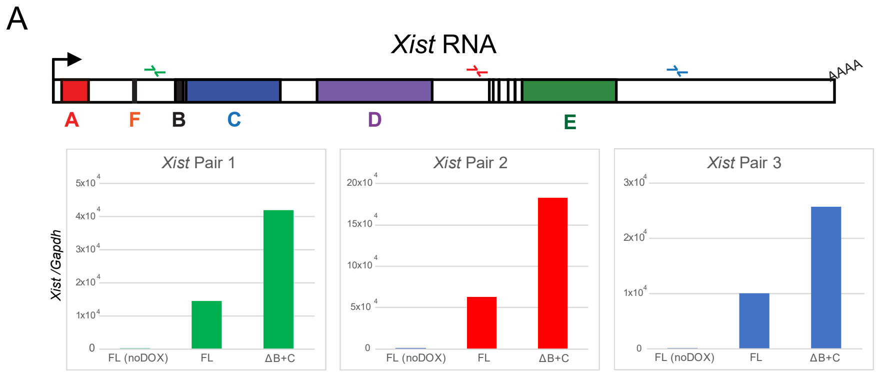


B

| Samples | % of cells with ***Xist*** **domains** |
|---|---|
| *Xist* FL noDOX | 0% |
| *Xist* FL DOX | 24.0% |
| *Xist* ΔB+C DOX | 50.9% |

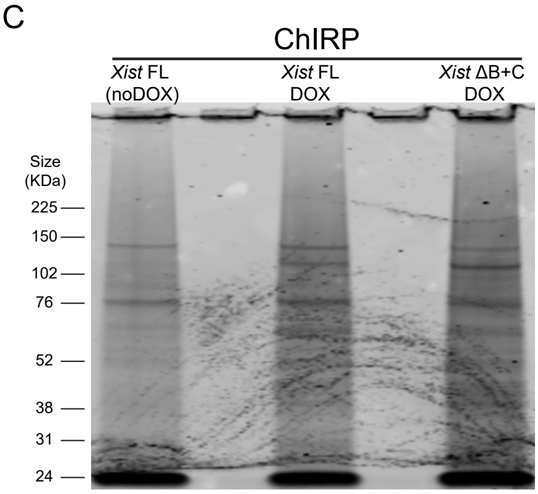


Figure 2 - figure supplement 1

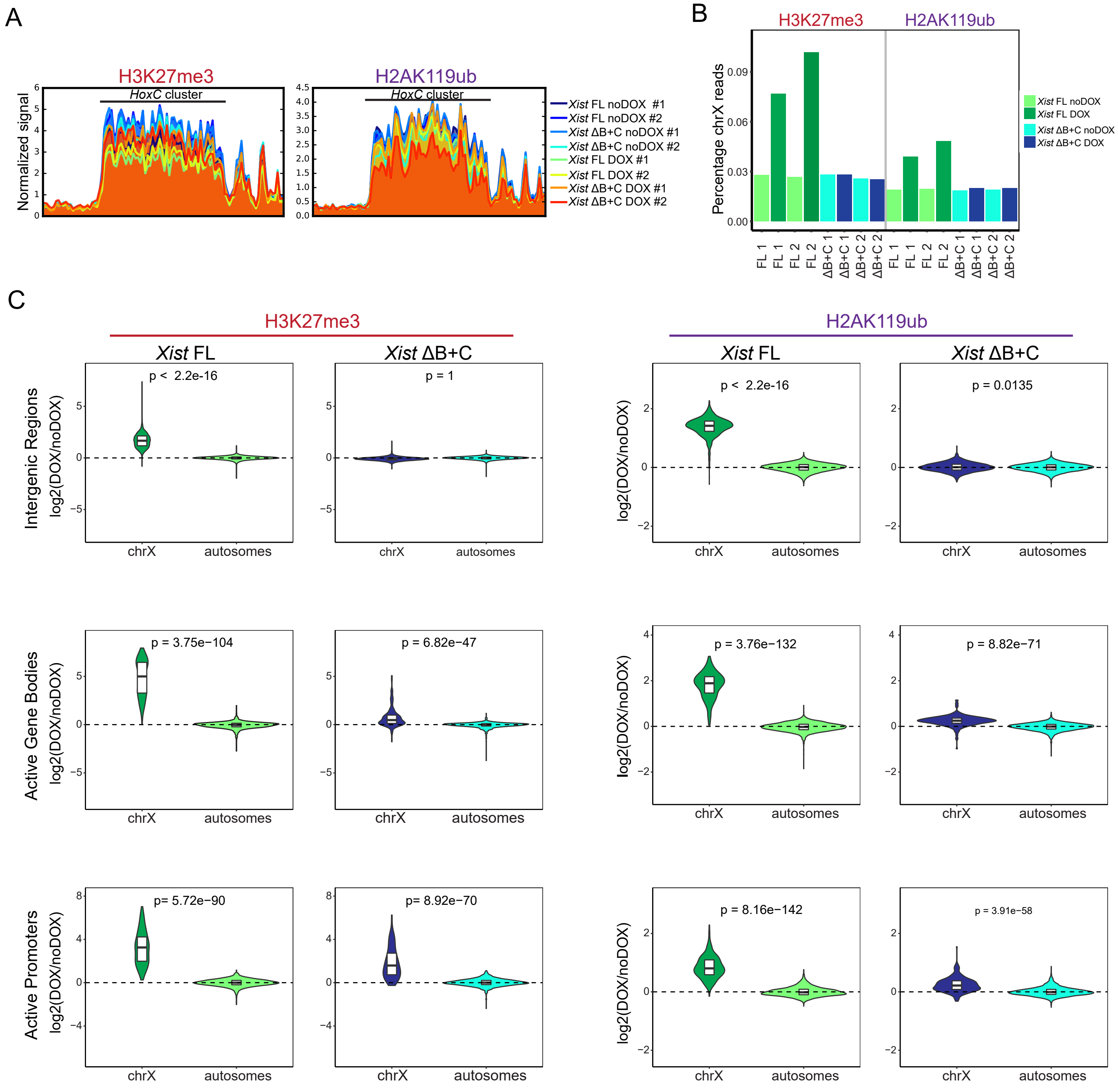


Figure 3 - figure supplement 1

A

| Samples | % of cells with ***Xist* domains** |
|---|---|
| *Xist* FL noDOX #1 | 0% |
| *Xist* FL noDOX #2 | 0% |
| *Xist* FL DOX #1 | 46.6% |
| *Xist* FL DOX #2 | 59.6% |
| *Xist* ΔB+C noDOX #1 | 0% |
| *Xist* ΔB+C noDOX #2 | 0% |
| *Xist* ΔB+C DOX #1 | 66.3% |
| *Xist* ΔB+C DOX #2 | 56.3% |

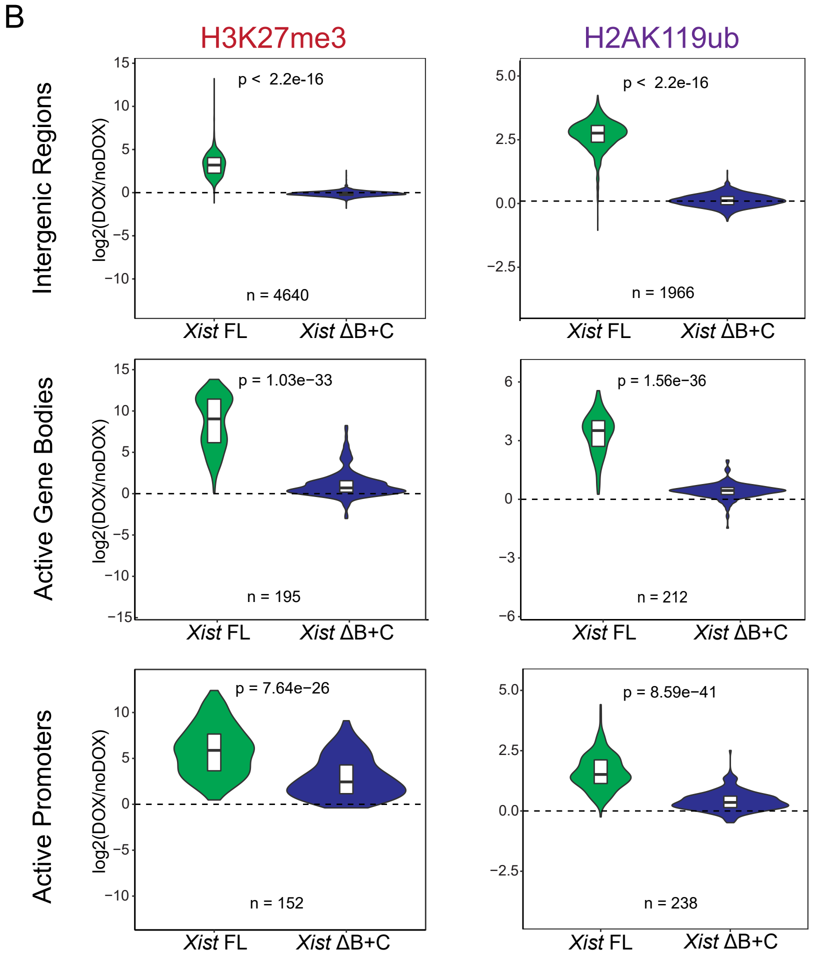


Figure 3 - figure supplement 2

A

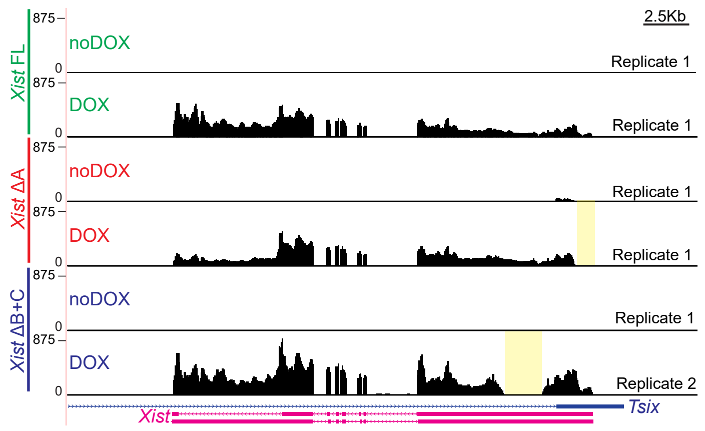


B

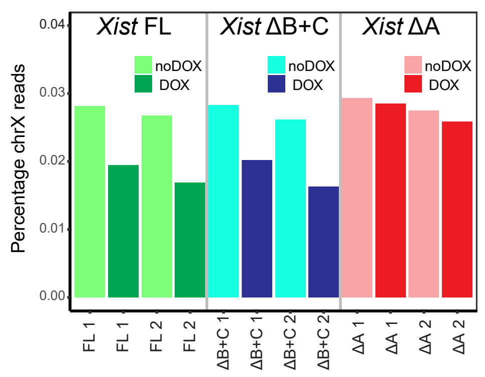


C

| Samples | Cells with *Xist*-coated chrX |
|---|---|
| *Xist* FL noDOX #1 | 0.00% |
| *Xist* FL noDOX #2 | 0.00% |
| *Xist* FL DOX #1 | 46.83% |
| *Xist* FL DOX #2 | 49.35% |
| *Xist* ΔA noDOX #1 | 0.00% |
| *Xist* ΔA noDOX #2 | 0.00% |
| *Xist* ΔA DOX #1 | 50.98% |
| *Xist* ΔA DOX #2 | 54.03% |
| *Xist* ΔB+C noDOX #1 | 0.00% |
| *Xist* ΔB+C noDOX #2 | 0.00% |
| *Xist* ΔB+C DOX #1 | 55.38% |
| *Xist* ΔB+C DOX #2 | 56.69% |

F

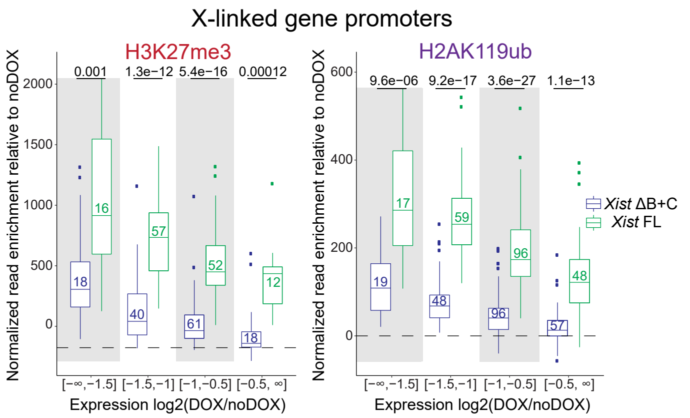


D

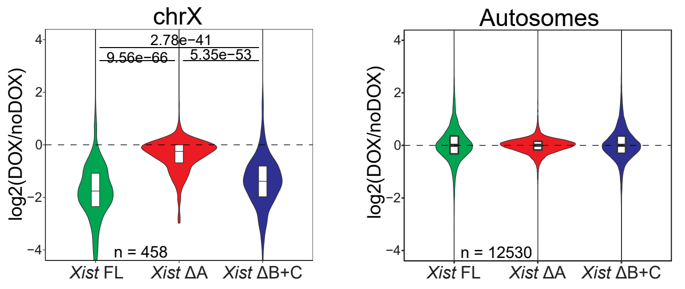


G

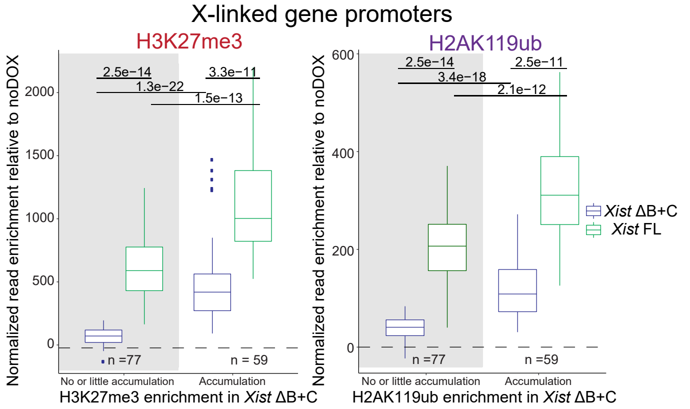


E

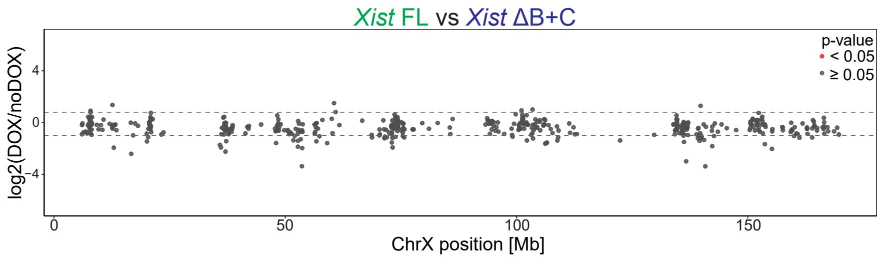


H

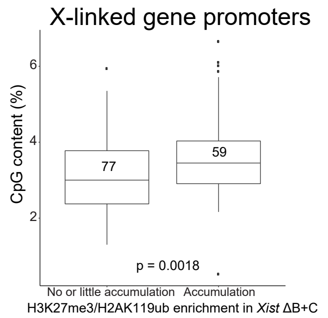


Figure 4 - figure supplement 1
